# PARP6-dependent vimentin ADP-ribosylation prevents myofibroblast activation in cardiac fibrosis

**DOI:** 10.64898/2026.02.18.706611

**Authors:** Sudharshana Sundaresan, Arushi Taneja, David Kubon, Apeksha Bhuyar, Anna Keodara, Deena M. Leslie Pedrioli, Arathi Bangalore Prabhashankar, Prasanna Simha Mohan Rao, Nagalingam Ravi Sundaresan, Michael O. Hottiger

## Abstract

Cardiac fibrosis is a central driver of adverse remodeling during heart failure, yet the post-translational regulation of myofibroblast activation remains poorly defined. Here, we identified PARP6 as a mono-ADP-ribosyltransferase that repressed myofibroblast activation through the ADP-ribosylation of vimentin. PARP6 expression was reduced in failing human hearts and *Parp6* haploinsufficiency in mice was sufficient to induce cardiac fibrosis. At the cellular level, PARP6 ADP-ribosylated vimentin thereby limiting actin stress fiber formation. Mechanistically, PARP6 inhibition enhanced RhoA activation and vimentin-RhoA complex formation, thus activating the RhoA-ROCK-LIMK-cofilin pathway. Consistently, Parp6 haploinsufficiency was associated with increased cofilin phosphorylation in mice hearts. In primary cardiac fibroblasts, PARP6 inhibition promoted RhoA-dependent actin stress fiber accumulation and induced myofibrotic protein expression. Together, these findings define a PARP6-vimentin(ADP-ribosylation)-RhoA axis that restrained contractility-driven fibroblast activation, indicating a cardioprotective role of PARP6 with potential therapeutic relevance for fibrotic heart disease.

## INTRODUCTION

Cardiac failure, the leading cause of global mortality, is the culmination of complex cellular and molecular events^1^. Studies indicate that most forms of cardiac failure are accompanied by varying degrees of cardiac fibrosis ^2^, a major cause for structural remodeling of the heart^2–4^. Cardiac remodeling is mediated by fibroblast proliferation, myofibroblast differentiation, contractility and invasion, and the deposition of extracellular matrix (ECM) proteins^3^. At the cellular level, these processes are accompanied by cytoskeletal changes, of which the assembly of monomeric G-actin into polymeric F-actin and the formation of actin stress fibers are particularly well characterized^5^. Actin stress fibers are central to fibroblast mechano-activation, in which increasing matrix stiffness and integrin engagement promote focal adhesion maturation^6^ and FAK/Src signaling^7,8^. Focal adhesion organization and ECM protein deposition are characteristic of myofibroblasts, which further reinforce actin polymerization^9^, ultimately leading to cardiac tissue remodeling and fibrosis. Recent evidence suggests that cardiac fibrosis can drive heart failure by promoting cardiomyocyte hypertrophy, apoptosis, and cardiac chamber dilation^10,11^.

In a failing fibrotic heart, the myocardial proteome is compromised by genetic mutations^12^, transcriptional^13^ and translational alterations^14–16^, and post-translational modifications (PTM)^17^. Although the roles of PTMs such as phosphorylation^17^ and acetylation^18^ in the cardiac proteome are well characterized, ADP-ribosylation has only recently emerged as a relevant PTM in the failing heart^19–21^. ADP-ribosylation is a PTM catalyzed by a family of 17 ADP-ribosyltransferases (ARTs), and occurs either as long, branched ADP-ribose polymers (PARylation) or as discrete single ADP-ribose units (MARylation)^22^. ARTs establish a direct functional link between the myocardial proteome and cellular energy metabolism by utilizing nicotinamide adenine dinucleotide (NAD^+^) as a substrate to catalyze ADP-ribosylation^23^. Since NAD^+^ availability is a rate-limiting factor for cardiac bioenergetic demands^24^, any imbalance in the NAD^+^/NADH redox potential promotes adverse tissue remodeling and the eventual progression to heart failure^23^. Notably, this relationship becomes particularly pathological during ART hyperactivation in cardiac stress, where excessive NAD^+^ consumption for ADP-ribosylation competes with cardiac energy demands, thereby exacerbating oxidative injury, impairing mitochondrial respiration, and promoting contractile dysfunction^23,25,26^. Thus, the intersection of NAD^+^ metabolism and ADP-ribosylation signaling highlights a critical vulnerability in the failing heart, underscoring the need to define the specific roles of ADP-ribosylation in cardiovascular pathophysiology.

Most current research in the field focuses on PARP1-mediated PARylation in cardiomyocytes, where it is primarily implicated in ischemia-reperfusion injury and heart failure^27^. Hyperactive PARP1 depletes cellular NAD⁺ pools, exacerbates oxidative stress, and promotes cardiomyocyte death^25^. Beyond ischemic injury, PARP1 activation promotes nuclear accumulation of the transcription factor FoxO3a and enhances the expression of autophagy-related genes in cardiomyocytes^25^. Conversely, PARP1 also exhibits cardioprotective effects by PARylating HSP90AB1, thereby indirectly activating cell-cycle kinases and promoting cardiomyocyte proliferation and cardiac regeneration^21^. This duality establishes ADP-ribosylation as a context-specific, essential yet under-characterized regulatory layer in cardiac biology. Beyond the PARP1-centric understanding of ADP-ribosylation in the heart, the role of MARylation remains poorly defined despite the fact that most ART family members predominantly catalyze MARylation^22^.

Previous studies identified PARP6 as a mono-ART with context-dependent roles in cancer^28,29^. PARP6 acts as a tumor suppressor in colorectal cancer by downregulating survivin expression, whereas it functions as an oncogene in gastric cancer by positively regulating survivin expression^28,29^. In breast cancer, PARP6 regulates centrosome integrity by MARylating Chk1 and reducing tumor cytotoxicity^30^. PARP6 is also catalytically implicated in dendrite morphogenesis and neuronal development, and mutations in PARP6 have been linked to neurodevelopmental disorders^31,32^. Beyond cancer and neurogenesis, PARP6 remains largely unexplored in cardiovascular biology, where fibroblast-driven cytoskeletal remodeling and ECM deposition are central determinants of maladaptive cardiac remodeling^10^. Furthermore, the role of PARP6 in fibroblast activation and fibrotic signaling remains unknown despite the emerging view that mono-ARTs act as stress-sensing signaling effectors^22,33^. Defining PARP6-dependent MARylation pathways in fibroblasts may therefore uncover a previously unrecognized link between metabolic stress, mechanoactivation, and cardiac remodeling that extends beyond the prevailing cardiomyocyte-centric ADP-ribosylation paradigm in heart failure. This gap provides the rationale for the present study, which examines how PARP6-mediated ADP-ribosylation contributes to F-actin remodeling, fibroblast activation, and ultimately cardiac fibrosis.

## RESULTS

### Expression of PARP6 is downregulated in failing human hearts

To identify the potential role of mono-ARTs in cardiac pathophysiology, we quantified the mRNA expression levels of different ARTs, including PARP3, PARP6, PARP8, PARP9 and PARP10 in the failing human myocardium. Of the analyzed mono-ARTs, we found that PARP6 mRNA levels were significantly downregulated in failing heart tissues compared to non-failing controls (Fig. 1a,b). There was also a marked decrease in PARP6 protein expression in these failing human hearts (Fig. 1c,d). Notably, these failing hearts exhibited extensive fibrosis (Fig. 1e), suggesting a potential link between PARP6 deficiency and the progression of heart failure.

**Fig. 1:**
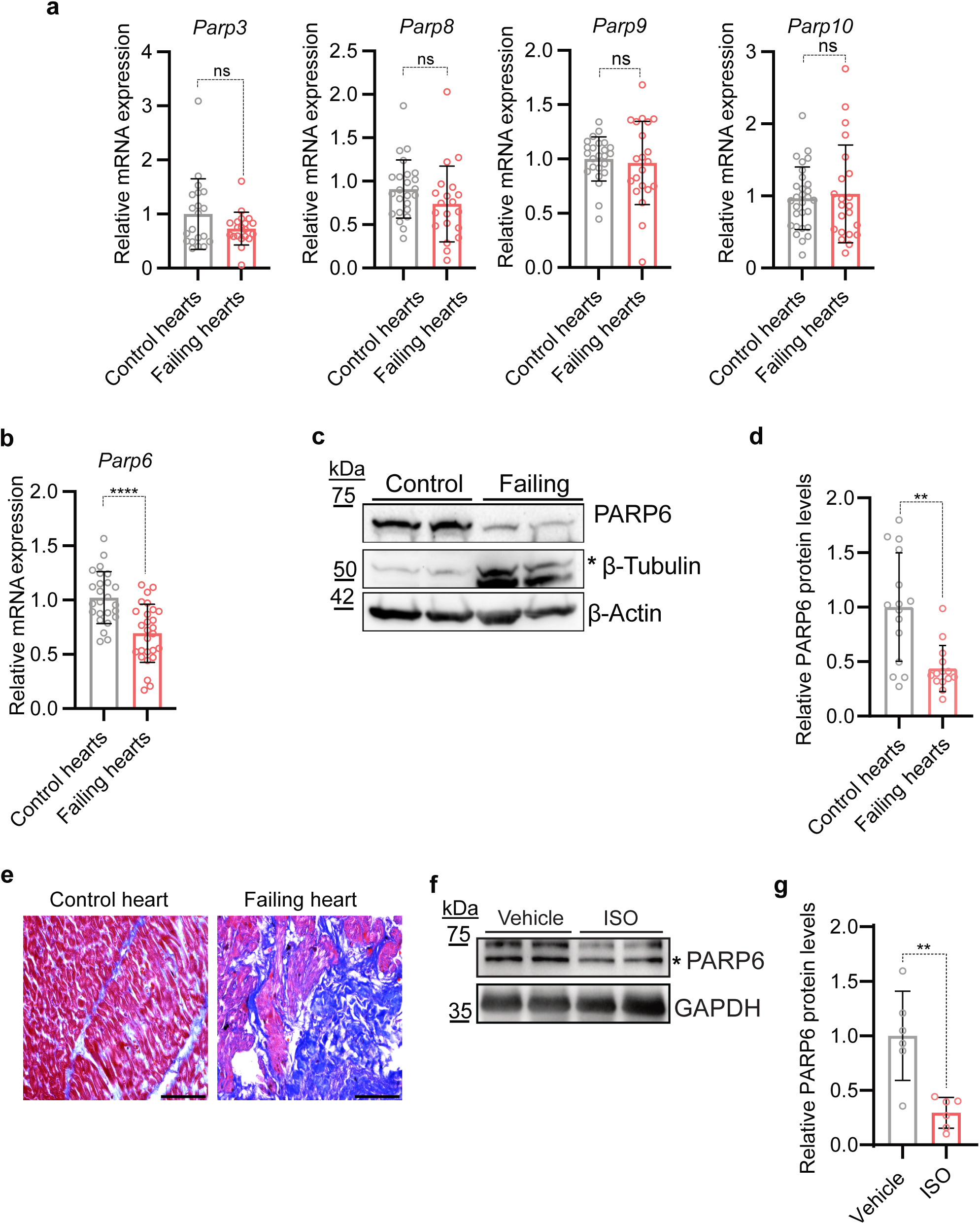
PARP6 is downregulated in failing human hearts. **a-b,** Relative mRNA expression of mono-ARTs in failing human hearts compared to healthy controls: **a**, Bar graphs show RNA expression of *Parp3, Parp8, Parp9, Parp10*, and **b,** *Parp6* in the indicated groups. Each bar represents group mean with error bars as plotted; statistical significance tests were determined using data normality analysis and performed by unpaired two-tailed Student’s t-tests or Man Whitney test. **c,** Representative immunoblot of PARP6 in healthy human heart samples and failing heart tissue lysates. β-actin and β-tubulin (band indicated by an asterisk) are shown as loading controls (n=15). **d,** Bar graph depicting the densitometric quantification of PARP6 immunoblot signal from **(c)**, normalized to β-Actin as loading control (n=14-15). Statistical significance was assessed by Man Whitney test. **e,** Representative Masson’s trichrome staining for control and failing human heart tissue samples. Scale bar represents 75 µm. **f,** Representative immunoblot of PARP6 (band indicated by an asterisk) in isoproterenol-infused mice hearts (ISO) compared to vehicle-infused control hearts (n=6). GAPDH is used as loading control. **g,** Densitometric quantification of (**f**). ns=not significant, * = p < 0.05, ** = p < 0.01, *** = p < 0.001, **** = p < 0.0001.

To further validate the role of PARP6 in the heart, we investigated a murine model of isoproterenol (ISO)-induced cardiac dysfunction^18^. In line with the failing human hearts, these ISO-infused hearts also exhibited a significant reduction in PARP6 protein expression (Fig. 1f, g), suggesting that PARP6 loss is a fundamental event in the pathogenesis of cardiac dysfunction.

### PARP6 haploinsufficiency promotes cardiac dysfunction and pathological tissue remodeling

To address whether the reduction in PARP6 levels is the cause or consequence of a failing myocardium, we generated a global *Parp6* heterozygous knockout mouse model using the Cre-*loxP* recombination system (Fig. 2a and Fig. S1a). We validated the monoallelic deletion of *Parp6* in mRNA and protein levels across multiple organs (Fig. S1b-g) and the reduction of PARP6 protein levels in the heart of 6-7-month-old *Parp6^+/−^* mice (Fig. 2b).

**Fig. 2:**
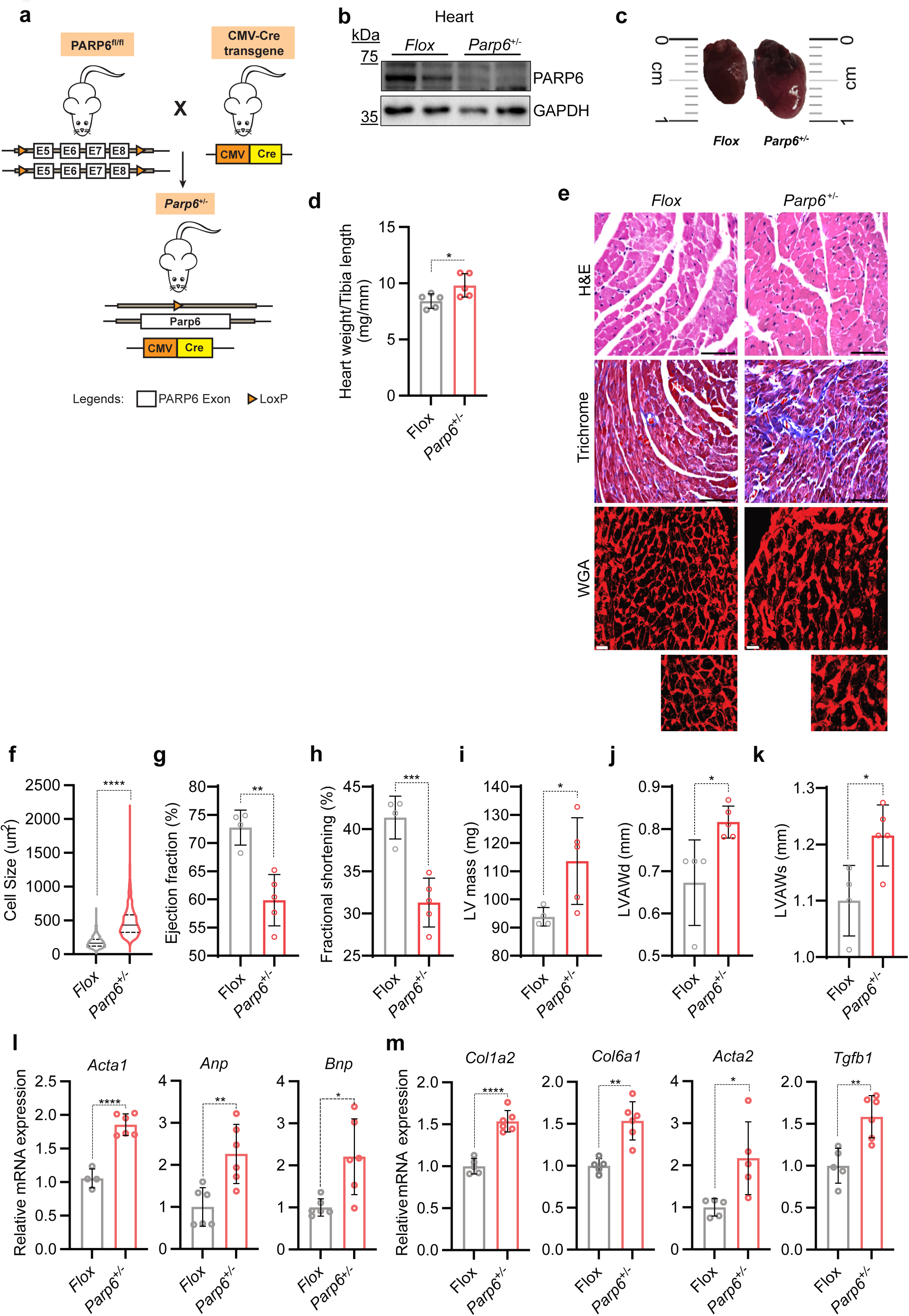
PARP6 haploinsufficiency promotes pathological remodeling and cardiac dysfunction. **a,** Schematic of the genetic targeting and breeding strategy for the generation of *Parp6*^+/−^ animals. **b,** Phenotype validation by immunoblot of PARP6 protein in heart lysates from Cre-negative *Parp6*^fl/fl^ (Flox) and *Parp6*^+/−^ mice; GAPDH is shown as loading control, (n=3). **c,** Comparative gross morphological representation of 6-7-month-old *Parp6*^+/−^ and age-matched Cre-negative control hearts (Flox). **d,** Bar graph quantifying the heart weight to tibia length ratio of Flox and *Parp6*^+/−^ mice. **e,** Representative hematoxylin/eosin (top panel; Scale bar represents 75µm), Masson’s trichrome stain (middle panel; scale bar represents 75µm) and wheat germ agglutinin staining (bottom panel scale bar represents 30µm) of left ventricular myocardium comparing Flox and *Parp6*^+/−^ hearts (overview and higher-magnification insets as indicated). **f,** Bar graphs showing the quantification of cardiomyocyte cross-sectional area (cell size); **g-k,** Echocardiographic parameters in 6-7-month-old *Parp6*^+/−^ and age-matched Cre controls mice: **g**, ejection fraction (%); **h,** fractional shortening (%); **i,** left ventricle (LV) mass; **j,** left ventricle anterior wall thickness at diastole (LVAWd), and **k,** left ventricle anterior wall thickness at systole (LVAWs) in Flox and *Parp6*^+/−^ hearts. **l-m,** Cardiac gene expression analysis by RT-qPCR. Bar graphs showing the relative mRNA expression of remodeling/stress-associated markers *Acta1, Anp, Bnp*; and **m,** extracellular matrix and pro-fibrotic markers *Col1a2, Col6a1, Acta2*, and *Tgfb1* in Flox and *Parp6*^+/−^ mouse hearts. Data are shown as relative expression (normalized to the indicated reference gene) with individual biological replicates overlaid; statistical significance was assessed by Student’s t-test (unpaired) or Man Whitney test; annotated on plots as ns (not significant), * p < 0.05, ** p < 0.01, *** p < 0.001, **** p < 0.0001.

To understand the cardiac-specific response to a single-copy deletion of *Parp6*, we measured the hearts of 6-7-month-old male *Parp6^+/−^*mice relative to *Parp6^fl/fl^* control animals. We observed a significant increase in the gross heart size in *Parp6^+/−^* mice, accompanied by an increase in their heart weight-to-tibia length ratio, indicative of cardiac hypertrophy (Fig. 2c, d). Histological analysis revealed increased interstitial fibrosis in *Parp6*^+/−^ mice hearts (Fig. 2e). Moreover, these hearts showed an increase in individual cardiomyocyte cross-sectional area (Fig. 2e,f). We also observed a significant increase in lung and spleen weights of *Parp6^+/−^* mice relative to controls (data not shown). Echocardiography analysis of these mice revealed contractile dysfunction, as evidenced by reduced ejection fraction and fractional shortening (Fig. 2g,h). These mice also showed increased overall LV mass, along with increased anterior wall diameters, consistent with overall heart enlargement (Fig. 2i-k). We observed an increase in the posterior wall diameters (non-significant), with no discernible changes in the internal diameters (Figure S1h-k). Cardiac dysfunction in *Parp6*^+/−^ mice was accompanied by activation of the fetal gene program, including skeletal muscle alpha-actin (*Acta1*), Atrial natriuretic peptide (*Anp*), and Brain natriuretic peptide (*Bnp*) (Fig. 2l).

Consistent with pathological remodeling, we sought to define the molecular drivers of fibrotic transformation at the transcript level. Upon assessment of a panel of profibrotic markers, we observed a marked increase in mRNA levels of *Col1A*, *Col6A, Acta2* (α-SMA), and *Tgfb1* (Fig. 2m), indicative of the activation of a fibrotic gene program^34^ in *Parp6^+/−^* mice.

### PARP6 ADP-ribosylates cysteine residues of cytoskeletal and cytoskeleton-regulating proteins in AC16 cardiomyoblasts

PARP6 is a 630-amino acid cytosolic mono-ART with an N-terminal RWD domain, a central cysteine-rich C4 zinc-finger domain, and a helical subdomain (HE) preceding the C-terminal ART catalytic domain (ART), which contains a conserved catalytic triad^35^ (Fig. S2a). To investigate the enzymatic function of PARP6 in cardiac cells, we first generated a doxycycline (Dox)-inducible overexpression model of HA-tagged wild-type PARP6 (PARP6^WT^) and its catalytically inactive mutant (PARP6^Y508A^) by mutating the conserved Tyr508 residue in its catalytic triad (Fig. S2a). Western blot analysis confirmed robust expression of both PARP6^WT^ and PARP6^Y508A^ constructs in the AC16 cardiomyocyte-fibroblast fusion cell line, with an increase in ADP-ribosylation signal only upon PARP6^WT^ overexpression, thereby validating PARP6^Y508A^ as a catalytically inactive mutant of PARP6^WT^ (Fig. S2b).

Although PARP6 has been reported to catalyze MARylation^36^, its comprehensive ADP-ribosylome and amino acid acceptor selectivity remain undefined. We, therefore, performed our in-house mass spectrometric ADP-ribosylome workflow using the macrodomain Af1521 and its engineered variant eAf1521 for the enrichment of ADP-ribosylated peptides and identified site-localized ADP-ribose by MS/MS^37^ (Fig. 3a). PARP6^WT^ overexpression increased ADP-ribose peptide-spectrum matches (PSM), unique ADP-ribose peptides, modified proteins, and ADP-ribose acceptor sites (Fig. 3b). Site localization analysis of amino acid acceptor residues further revealed that PARP6^WT^ predominantly modified proteins at cysteine residues, establishing PARP6 as a writer of cysteine ADP-ribosylation (Fig. 3c). Notably, PARP6^WT^ was automodified at Cys215, further confirming its activity as a writer of cysteine ADP-ribosylation (Fig. S2c). Compartmental annotation of ADP-ribosylated proteins revealed an enrichment of cysteine-modified PARP6 target proteins in the cytoskeletal and cytoplasmic compartments with additional nuclear targets (Fig. 3d). Consistent with this observation, STRING analysis of ADP-ribosylated proteins with ≥1.5-fold higher spectral counts in PARP6^WT^ overexpression than in PARP6^Y508A^ overexpression revealed a densely connected cytoskeletal and cytoskeleton-regulating protein network (Fig. 3e). Furthermore, gene ontology (GO) analysis showed that the targets of PARP6 were enriched for cytoskeletal organization, thereby establishing PARP6 as a preferential writer of cytoskeletal and cytoskeleton-regulating proteins (Fig. 3f). In contrast, fewer proteins were preferentially modified in PARP6^Y508A^ overexpression than in PARP6^WT^ overexpression, which could indicate ADP-ribosylation mediated by either endogenous PARP6 or by other ARTs interacting with overexpressed PARP6^Y508A^ (Fig. S2d). However, no statistically significant GO enrichment terms were obtained after multiple-testing correction for the proteins preferentially modified in PARP6^Y508A^ overexpression (Fig. S2e), supporting the conclusion that PARP6^WT^ catalyzes a target-specific, cysteine-biased cytoskeletal ADP-ribosylation in AC16 cells.

**Fig. 3:**
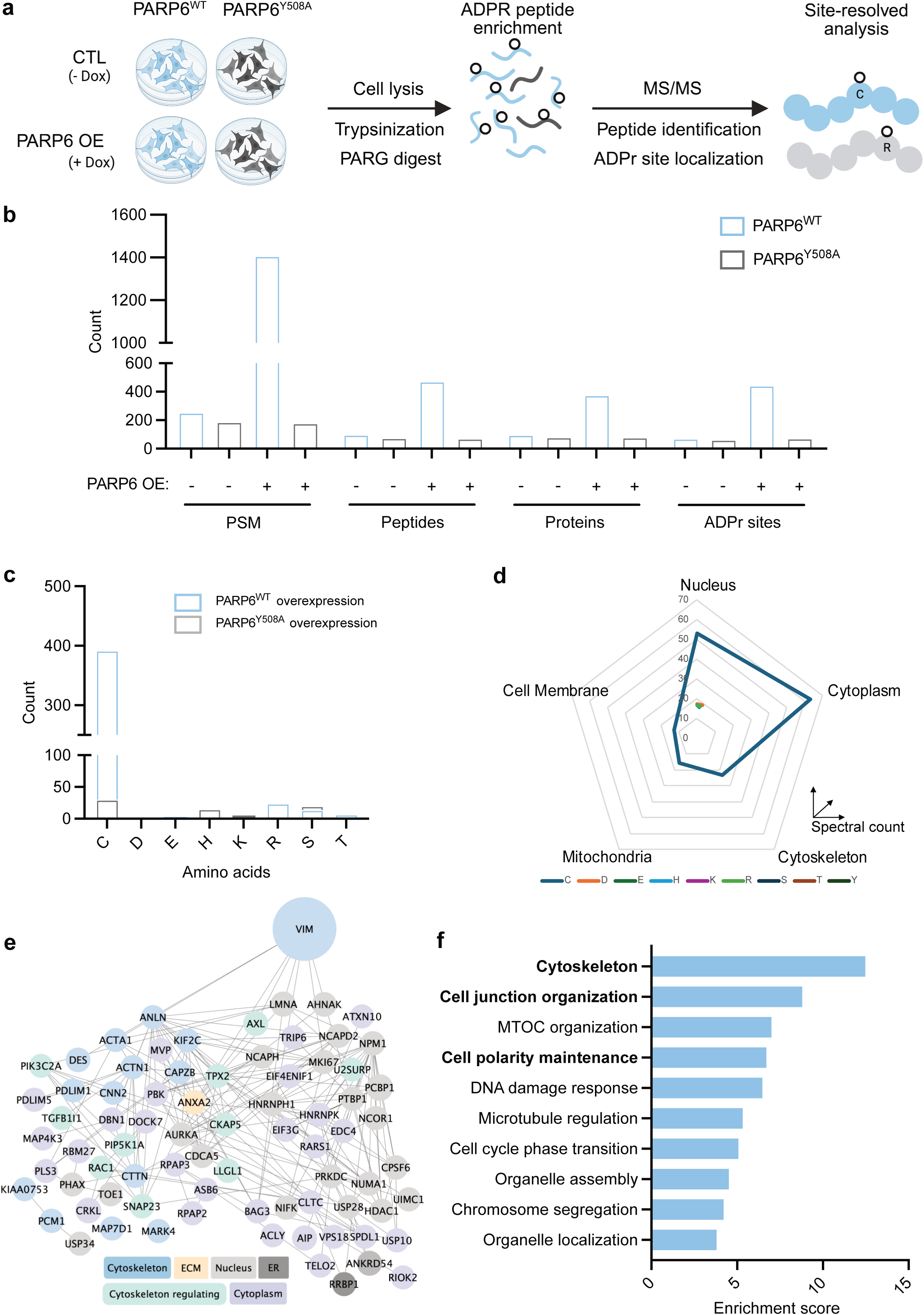
PARP6 ADP-ribosylates cysteine residues of cytoskeletal and cytoskeleton-regulating proteins in AC16 cardiomyoblasts. **a,** Schematic of the MS-based ADP-ribosylome workflow of PARP6^WT^ and PARP6^Y508A^ overexpression in AC16 cardiomyoblasts (induced with 250ng/mL doxycycline).**b,** Bar graph depicting the summary metrics (peptide-spectrum matches (PSMs), unique ADPr peptides, proteins, and localised ADPr sites) across PARP6^WT^ and PARP6^Y508A^ overexpression conditions. **c,** Overlaid bar chart depicting the amino-acid acceptor residues of PARP6^WT^ vs PARP6^Y508A^ overexpression conditions (single letter amino acid codes: C, D, E, H, K, R, S, T, Y; residues with zero PSM omitted). **d,** Radar plot of ADPR spectral counts across annotated cellular compartments (concentric gridlines in 10-count increments) across acceptor amino acids upon PARP6^WT^ overexpression conditions. **e,** STRING interaction network of proteins with ≥1.5-fold higher ADPr spectral counts in PARP6^WT^ overexpression compared to PARP6^Y508A^ overexpression; node size scaled with ADPr spectral counts, and node colour encodes depicted subcellular localisation. **f,** Gene Ontology-term enrichment (FDR < 0.05) for proteins with ≥1.5-fold higher ADPr spectral counts in PARP6^WT^ overexpression compared to PARP6^Y508A^ overexpression.

### PARP6 preferentially modifies vimentin at Cys328 while preserving its structural properties

We identified vimentin, a type III intermediate filament protein highly expressed in fibroblasts^38^, as the most abundant target of PARP6 (Fig. 3e). Among the multiple modification sites on vimentin, we identified Cys328 as the most modified residue, closely followed by Arg207 and, to a much lesser extent, by Arg196 (Fig. 4a). All type III intermediate filament proteins, including vimentin, desmin, glial fibrillary acidic protein, and peripherin, possess a cysteine residue that is conserved across species and, except in peripherin, is the only cysteine residue of the protein^39^. To confirm that vimentin ADP-ribosylation is PARP6-dependent, we validated AZ0108^40^ as a pharmacological PARP6 inhibitor (PARP6i) (Fig. S3a). Since AZ0108 has been reported to inhibit PARP1/2^41^, we used talazoparib, a clinically validated PARP1/2 inhibitor^42^ (PARP1/2i), to distinguish between the effects of PARP6 inhibition from those of PARP1/2 inhibition. We then immunoprecipitated vimentin from AC16 cells under PARP6 inhibition, PARP6^WT^, and PARP6^Y508A^ overexpression following washing with 1,6-hexanediol to eliminate unspecific hydrophobic interactions^43^ characteristic of vimentin filaments (Fig. 4b). We observed a reduction in vimentin ADP-ribosylation upon PARP6 inhibition, and an increase in vimentin ADP-ribosylation upon PARP6^WT^ overexpression, confirming that vimentin ADP-ribosylation is driven by the catalytic activity of PARP6 (Fig. 4b).

**Fig. 4:**
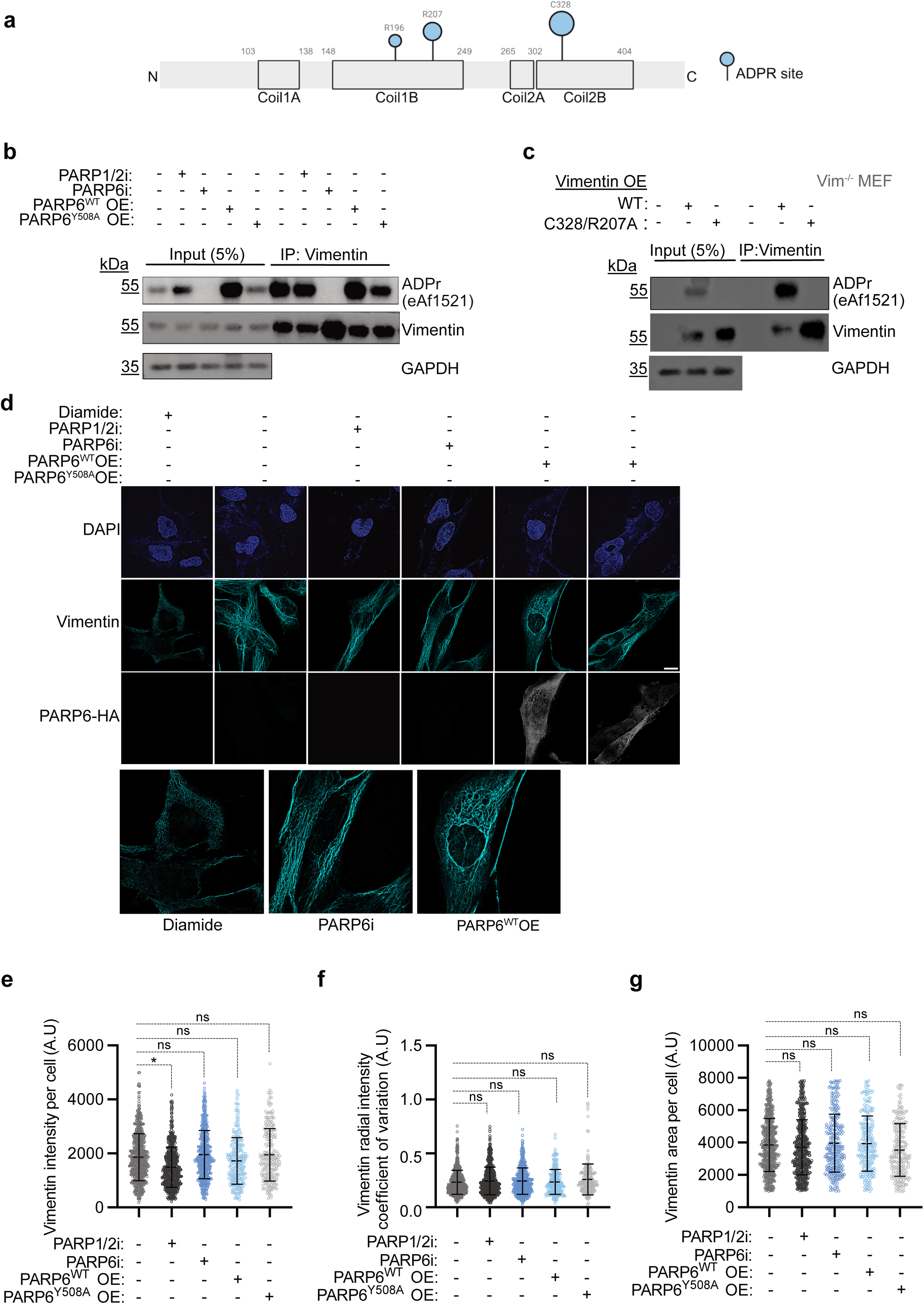
PARP6 preferentially modifies vimentin at Cys328 while preserving its structural properties. **a,** Domain schematic of human vimentin indicating coiled-coil subdomains (Coil 1A/1B/2A/2B) and the ADP-ribosylation sites detected (C328, R207, R196). The size of the modification symbol correlates with the spectral counts of modification on the marked acceptor site. **b,** Representative immunoblot of vimentin immunoprecipitation from AC16 cells under PARP1/2 inhibitor (PARP1/2i, 100 nM), PARP6 inhibitor (PARP6i, 1 µM), PARP6^WT^ or PARP6^Y508A^ overexpression (250 ng/mL doxycycline) conditions. GAPDH is shown as loading control (n=3). **c,** Representative immunoblot of vimentin immunoprecipitation from Vimentin^−/−^ MEF cells overexpressing vimentin^WT^ or vimentin^C328/R207A^. GAPDH is shown as loading control (n=2). **d,** Representative super-resolution images of AC16 cells under diamide (100 µM), PARP1/2i treatment (100 nM), PARP6i treatment (1µM), PARP6^WT^ overexpression and PARP6^Y508A^ overexpression (induced with 250 ng/ml doxycycline) conditions. Images show DAPI (nuclei, blue), vimentin (cyan), and PARP6-HA (grey). The bottom row shows magnified insets corresponding to the indicated conditions (n=3). Scale bar represents 20 μm. **e-g,** Scatter plots depicting quantifications of (**d**): **e,** vimentin fluorescence intensity per cell (A.U); **f,** vimentin radial intensity coefficient of variation per cell (A.U.); and **g,** vimentin area per cell. Scatter plots display single-cell distributions, and at least 200 cells were quantified per condition using confocal microscopy. Statistical testing was performed on the means of 3 independent experiments using one-way ANOVA with multiple-comparison correction. ns=not significant; * = p < 0.05; **** = p < 0.0001.

To validate that Cys328 and Arg207 are crucial for vimentin ADP-ribosylation, we generated vimentin C328A and R207A point mutants, as well as a combined C328/R207A mutant, and expressed these constructs in *Vim*^−/−^ mouse embryonic fibroblasts (MEFs) (Fig. S3b). Immunoprecipitation of vimentin from these cells revealed ADP-ribosylation of vimentin^WT^, while the vimentin^C328/R207A^ mutant exhibited no detectable ADP-ribosylation signal, establishing Cys328 and Arg207 as the primary acceptor sites of PARP6-dependent vimentin ADP-ribosylation (Fig. 4c). Consistent with previous studies of vimentin Cys328 mutants^44^, the C328A mutant did not form filaments but rather punctate foci in *Vim*^−/−^ MEFs. In contrast, the R207A mutant exhibited a ring-like perinuclear localization, which is uncharacteristic of vimentin^WT^ (Fig S3b).

Given that Cys328 of vimentin is a known hotspot for electrophilic PTMs that influence filament remodeling^39,44^, we tested whether PARP6-dependent vimentin ADP-ribosylation perturbs vimentin filament architecture. Super-resolution imaging showed an intact, extended vimentin network across PARP6i, PARP1/2i, PARP6^WT,^ and PARP6^Y508A^ overexpressing conditions in both AC16 cardiomyoblasts and primary cardiac fibroblasts, suggesting that vimentin filament formation was not affected by ADP-ribosylation (Fig. 4d and Fig. S3c) but could be abrogated by Cys oxidation with diamide (Fig. 4d). Quantifications of vimentin intensity, radial intensity heterogeneity (coefficient of variation of vimentin intensity from cell center to periphery; higher value reflects greater spatial unevenness), and area covered per cell number revealed no significant differences, thereby suggesting no detectable effect on bulk filament organization (Fig. 4e-g). This indicated that the downstream impacts of PARP6 perturbation were not attributable to ADP-ribosylation-mediated structural collapse of vimentin.

Furthermore, since Cys328 of vimentin has been linked to regulating lysosomal positioning^44,45^, we assessed LAMP1 intensity and distribution after PARP6i treatment and after PARP6 overexpression (Fig. S3d,e). Lysosomal distribution was unchanged between control, PARP1/2i, and PARP6i treatments as assessed by LAMP1 staining, with modest changes across both PARP6^WT^ and PARP6^Y508A^ overexpression, indicating overexpression-related cellular stress rather than an effect of PARP6-dependent vimentin ADP-ribosylation (Fig. S3d,e).

### PARP6-mediated vimentin ADP-ribosylation attenuates actin stress fiber formation

Intermediate filaments coordinate with both actin microfilaments^46^ and microtubules^47^ to form an integrated cytoskeletal network that regulates cellular responses to biochemical and mechanical cues^48^. In particular, the bidirectional relationship between the vimentin and actin networks has recently gained importance, with vimentin characterized as interacting with actin both directly^49^ and through the linker protein plectin^50^ to regulate actin organization during the cell cycle^49^ and under oxidative stress^51^. We thus investigated whether PARP6-mediated vimentin ADP-ribosylation could regulate actin network remodeling and actin remodeling-induced myofibroblast activation^9^. To define the contribution of vimentin to actin reorganization in AC16 cardiomyoblasts, we depleted vimentin from AC16 cardiomyoblasts and observed a significant increase in F-actin stress fibers throughout the cytoplasm (Fig. 5a,b). The organization of actin stress fibers generates cellular traction forces that lead to focal adhesion-dependent changes in cell shape, area, and polarity^52^. Consistent with the increase in traction forces upon increased stress fiber organization, we observed a significant increase in the cell surface area of vimentin-knockdown cells (Fig. 5a,c). These observations identify vimentin as a cell-intrinsic repressor of actin stress fiber assembly in AC16 cells.

**Fig. 5:**
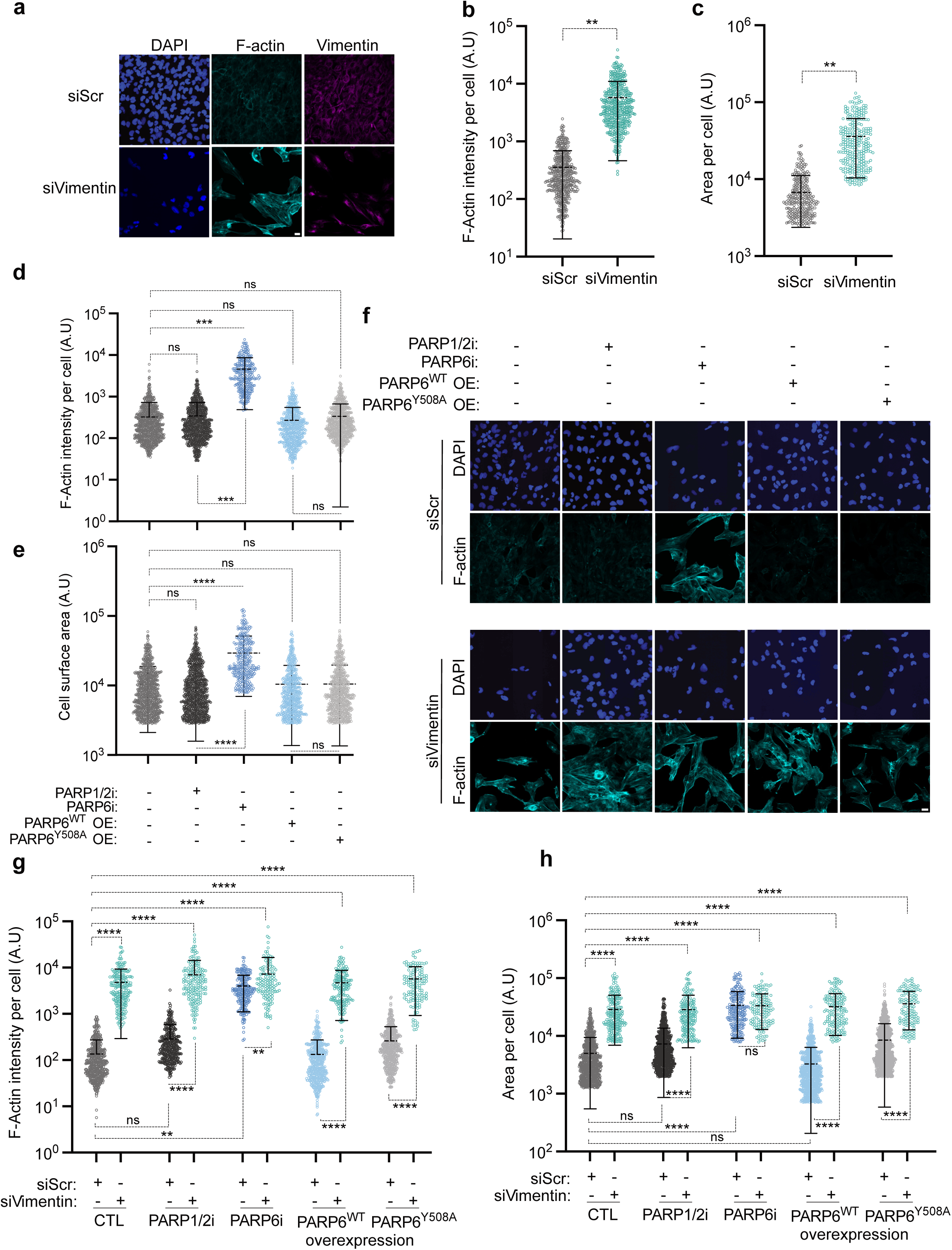
PARP6-mediated vimentin ADP-ribosylation attenuates actin stress fiber formation. **a,** Representative confocal microscopy images of scrambled siRNA (siScr) and siVimentin-treated cells. Images show DAPI (blue), F-actin (cyan), and vimentin (magenta) (n=3). Scale bar represents 20 μm. **b-c,** Scatter plots depicting quantifications of (**a**): **b**, F-actin intensity per cell; **c,** cell surface area per cell. At least 400 cells were quantified per condition. Statistical testing was performed on the means of 3 independent experiments using one-way ANOVA with multiple-comparison correction. **d-e,** Scatter plot quantification of immunofluorescence stainings under PARP1/2i (100 nM), PARP6i (1 µM) or PARP6 overexpression conditions (induced with 250 ng/ml doxycycline) from figure S3a; **d**, F-actin intensity per cell; **e,** cell surface area per cell (n=3; at least 300 cells quantified per condition). Statistical testing was performed on the means of 3 independent experiments using one-way ANOVA with multiple-comparison correction. **f,** Representative confocal microscopy images of siScr (10 nM) and siVimentin (10 nM) treated cells under PARP1/2i (100 nM), PARP6i (1 µM) or PARP6 overexpression conditions (induced by 250 ng/ml doxycycline). Images show DAPI (blue) and F-actin (cyan) (n=3) Scale bar represents 20 µm. **g-h,** Violin plot quantification of (**f**): **g**, F-actin intensity; **h,** cell surface area per cell (n=3; at least 140 cells quantified per condition). Statistical testing was performed on the means of 3 independent experiments using one-way ANOVA with multiple-comparison correction. Ns = non-significant, * = p < 0.05, ** = p < 0.01, *** = p < 0.001, **** = p < 0.0001.

To establish a role for PARP6-dependent vimentin ADP-ribosylation in actin stress fiber formation, we treated cells with PARP6i and found that PARP6i treatment phenocopied vimentin knockdown by significantly increasing F-actin stress fiber formation and cell surface area relative to control conditions (Fig. 5d,e and Fig. S4a). Overexpression of PARP6^WT^, but not of PARP6^Y508A^, showed a modest trend towards reduced baseline F-Actin stress fibers and cell surface area compared to control conditions, supporting the notion that PARP6 catalytic activity is required to maintain the low-stress-fiber state. (Fig. 5d,e and Fig. S5a). We did not reconfirm these findings using vimentin ADP-ribose-site mutants, as C328A substitution of vimentin disrupts filament assembly^46^, making it difficult to distinguish between the effects of loss of vimentin ADP-ribosylation and impaired scaffold integrity (Fig. S3b). Instead, we combined vimentin knockdown with PARP6 inhibition, PARP6^WT^, and PARP6^Y508A^ overexpression to understand if ADP-ribosylation of vimentin acts as the regulator of actin stress fiber formation (Fig. 5f). Under siScr conditions, PARP6 inhibition robustly increased F-actin stress fiber formation and cell spreading. In contrast, upon vimentin knockdown, PARP6 inhibition did not produce an additional increase in F-actin intensity or cell surface area compared to vimentin knockdown conditions alone (Fig. 5f-h). This observed phenotype was consistent with the model in which vimentin ADP-ribosylation restrained actin stress fiber organization in fibroblasts. Conversely, PARP6^WT^ overexpression failed to rescue the increased actin stress fiber formation upon vimentin depletion, indicating that PARP6 catalytic activity alone is insufficient to regulate actin stress fibers in the absence of vimentin as its substrate (Fig. 5f-h). Together, the non-additive nature of PARP6 inhibition and vimentin knockdown supported a model in which PARP6 functioned upstream of vimentin and constrained actin stress fiber assembly via vimentin ADP-ribosylation, rather than acting through a vimentin-independent mechanism.

### Proteomic analysis of PARP6 inhibition suggests Rho-GTPase activation and pro-fibrotic remodeling

Actin stress fiber assembly is not driven by G-actin alone, but by the coordinated interplay of actin-binding proteins that serve as filament nucleators, crosslinkers, and stabilizers, including myosin motors, focal-adhesion components that anchor actin bundles to integrins and transmit force, as well as the ECM, which mechano-modulates actin stress fiber assembly in a feedback loop^53^. To identify factors relevant to vimentin ADP-ribosylation-dependent regulation of actin stress fibers, we performed quantitative proteomics of PARP6i-treated cells. We identified multiple proteins exhibiting differential abundance upon PARP6i treatment compared to control conditions (Fig. 6a). STRING analysis of proteins with increased abundance under PARP6 inhibition revealed a highly connected network enriched for cell-matrix and tissue remodeling programs, which were reconfirmed by GO-term analysis (Fig. 6b,c). This suggested that PARP6 inhibition-mediated regulation of the actin cytoskeleton is accompanied by broader proteomic reprogramming that shifts cells toward a mechanically engaged, actin-remodeling-competent state. Conversely, proteins that decreased with PARP6 inhibition formed a less connected STRING network, whose GO term analysis was enriched for nuclear proteins and protein-DNA complexes (Fig. 6d,e).

**Fig. 6:**
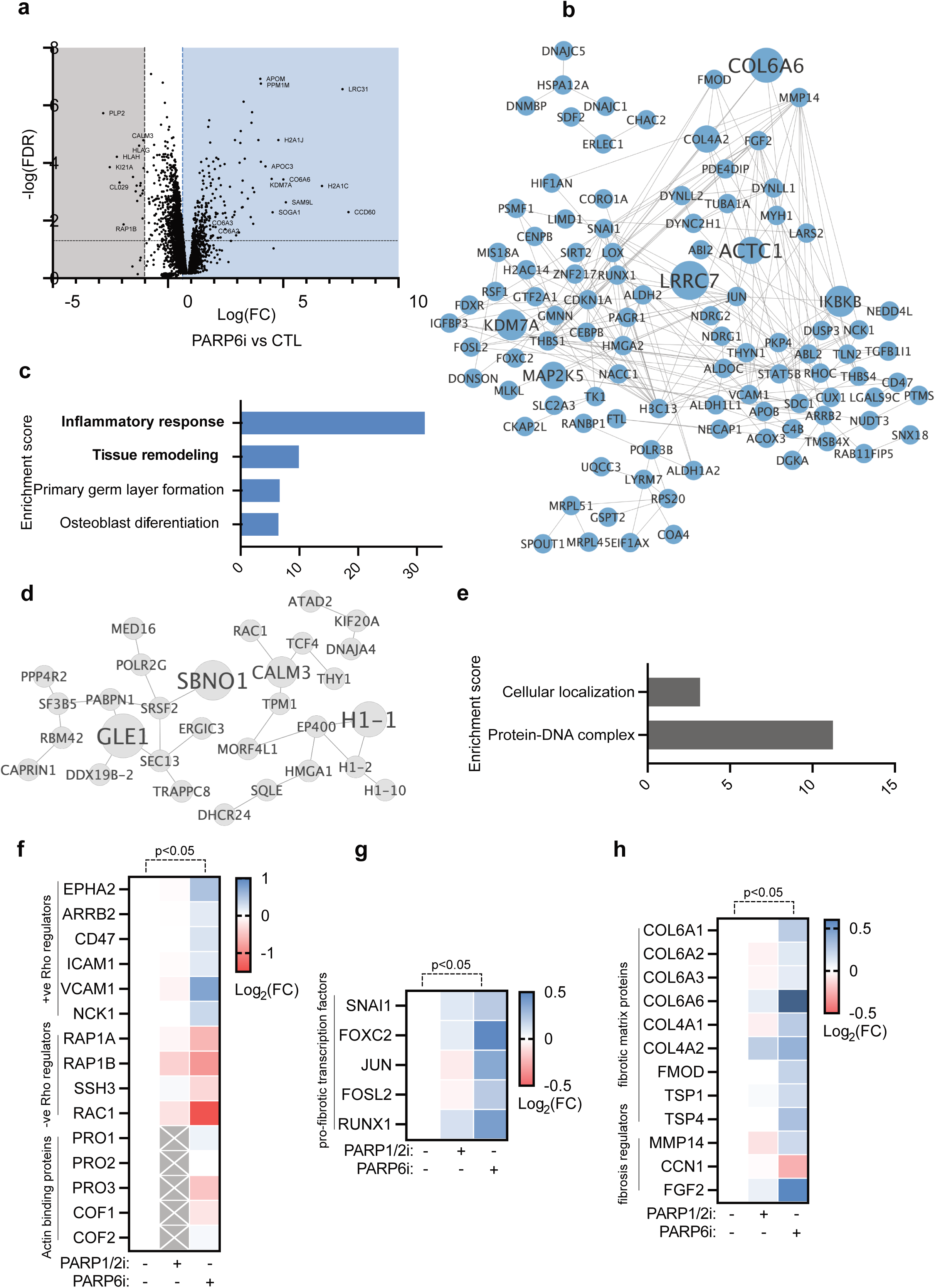
Proteomic analysis of PARP6 inhibition suggests Rho-GTPase activation and pro-fibrotic remodeling. **a,** Volcano plot depicting differential protein abundance in PARP6i treatment (1 µM) versus control conditions. Dashed lines indicate fold-change (1.5x in abundance increase, 0.5x in decrease) and FDR (q < 0.05) cutoffs. Significant proteins are highlighted (up in blue, down in grey) and select hits labelled (n=3). **b,** STRING interaction network of proteins with ≥1.5-fold higher abundance in PARP6i-treated conditions compared to control with FDR<0.05 as identified in **(a)**; node size scaled to fold change. Unconnected nodes were omitted. **c,** Gene Ontology-term enrichment (FDR < 0.05) for proteins with ≥1.5-fold higher abundance in PARP6i-treated conditions compared to control conditions as identified in **(a)**. **d,** STRING interaction network of proteins with ≤0.5-fold lower abundance in PARP6i-treated conditions compared to control with FDR<0.05 as identified in **(a)**; node size scaled to fold change. Unconnected nodes were omitted. **e,** Gene Ontology-term enrichment (FDR < 0.05) for proteins with ≤0.5-fold lower abundance in PARP6i-treated conditions compared to control conditions, as identified in **(a)**. **f-h,** Heatmaps showing abundance changes of **(f)** Rho-GTPase regulating proteins; **g,** pro-fibrotic transcription factors; and **h,** fibrotic proteins and regulators (FDR < 0.05) in PARP1/2i (100 nM) and PARP6i-treated (1 µM) conditions as identified in **(a).**

Interestingly, among the tissue remodeling proteins, PARP6 inhibition altered the abundance of several proteins previously implicated in regulating actin cytoskeletal remodeling, particularly of the regulators of Rho-GTPase signaling (Fig. 6f-h). Rho family GTPases are molecular switches that cycle between an inactive GDP-bound and an active GTP-bound state to coordinate actin cytoskeletal dynamics and mechanotransduction signaling^54^. They are modulated by guanine nucleotide exchange factors (GEFs) and GTPase-activating proteins (GAPs)^55^. Upon PARP6 inhibition, we found a significant increase in several Rho-GTPase activating proteins^56–61^, and a concurrent decrease in several Rho-GTPase repressing proteins^62,63^ (Fig. 6f). In parallel, PARP6 inhibition increased the abundance of several transcription factors associated with pro-fibrotic signaling or mesenchymal-reprogramming, (including AP-1 family components^64,65^ and SNAI1^66^) as well as extracellular matrix components and several fibrosis signaling regulators^67^, all of which can directly or indirectly mechanomodulate actin stress fiber formation through Rho-GTPase signaling (Fig. 6g,h).

### PARP6-dependent vimentin ADP-ribosylation prevents RhoA hyperactivation and restrains RhoA-ROCK-LIMK signaling

Among the Rho-GTPases, RhoA is the best-studied member, serving as the principal driver of actomyosin contractility and myofibroblast transition^68^. Furthermore, RhoA has previously been implicated in vimentin-dependent actomyosin contractility^51,69^. We therefore tested whether the ADP-ribosylation of vimentin is coupled to altered vimentin-RhoA complex formation and if the modified complex formation modulates RhoA activation state. Consistent with this model, immunoprecipitation of vimentin led to an increased co-immunoprecipitation of RhoA upon PARP6 inhibition, confirming that vimentin-RhoA complex formation is dependent on the ADP-ribosylation state of vimentin (Fig. 7a). To determine whether PARP6 inhibition alters the activation state of RhoA, we performed a rhotekin Rho-binding domain (RBD) pulldown to selectively enrich the GTP-bound active pool of RhoA (IP: RhoA^GTP^) (Fig. 7b). PARP6 inhibition increased RhoA^GTP^ abundance and concomitantly increased the amount of co-immunoprecipitated vimentin whereas the overexpression of PARP6^WT^ reduced RhoA^GTP^ levels (Fig. 7b), suggesting that PARP6-mediated vimentin ADP-ribosylation restrained RhoA activation and thereby limited the interaction of vimentin with RhoA^GTP^. RhoA can signal through multiple effector proteins, including ROCK I/II^6^, citron kinase^70^, myosin phosphatase^71^, and Dia^72^. To map the signaling downstream of RhoA, we first inhibited ROCK using Y-27632 (ROCKi) under PARP6 inhibition and vimentin knockdown conditions and confirmed the loss of actin stress fibers (Fig. S5a-d). This suggested that the actin stress fibers induced by PARP6i treatment and vimentin knockdown are ROCK-dependent (Fig. S5a-d). Upon RhoA-dependent activation, ROCK phosphorylates and activates LIMK^73^, which in turn phosphorylates cofilin^74^ at Ser3 to inhibit its actin-severing activity^75^. We therefore knocked down ROCK, LIMK, and cofilin under PARP6i treatment and vimentin knockdown conditions (Fig. 7c-g). Furthermore, since prior work has shown that vimentin can restrain GEF-H1-dependent RhoA activation in osteosarcoma cells^85^, we also knocked down GEF-H1, a potential upstream activator of RhoA (Fig. 7c-f). Under both vimentin depletion and PARP6 inhibition conditions, depletion of RhoA, ROCK, and LIMK strongly attenuated actin stress fiber induction, consistent with the canonical RhoA-ROCK-LIMK-cofilin signaling pathway (Fig. 7c-g). Knockdown of GEF-H1 did not rescue the induction of actin stress fibers, suggesting an alternate mechanism of RhoA activation in AC16 cardiomyoblasts (Fig. 7c-f). In line with this signaling, vimentin knockdown increased cofilin phosphorylation, confirming impaired actin disassembly and a consequent impaired actin stress fiber turnover upon PARP6 inhibition (Fig. 7g).

**Fig. 7:**
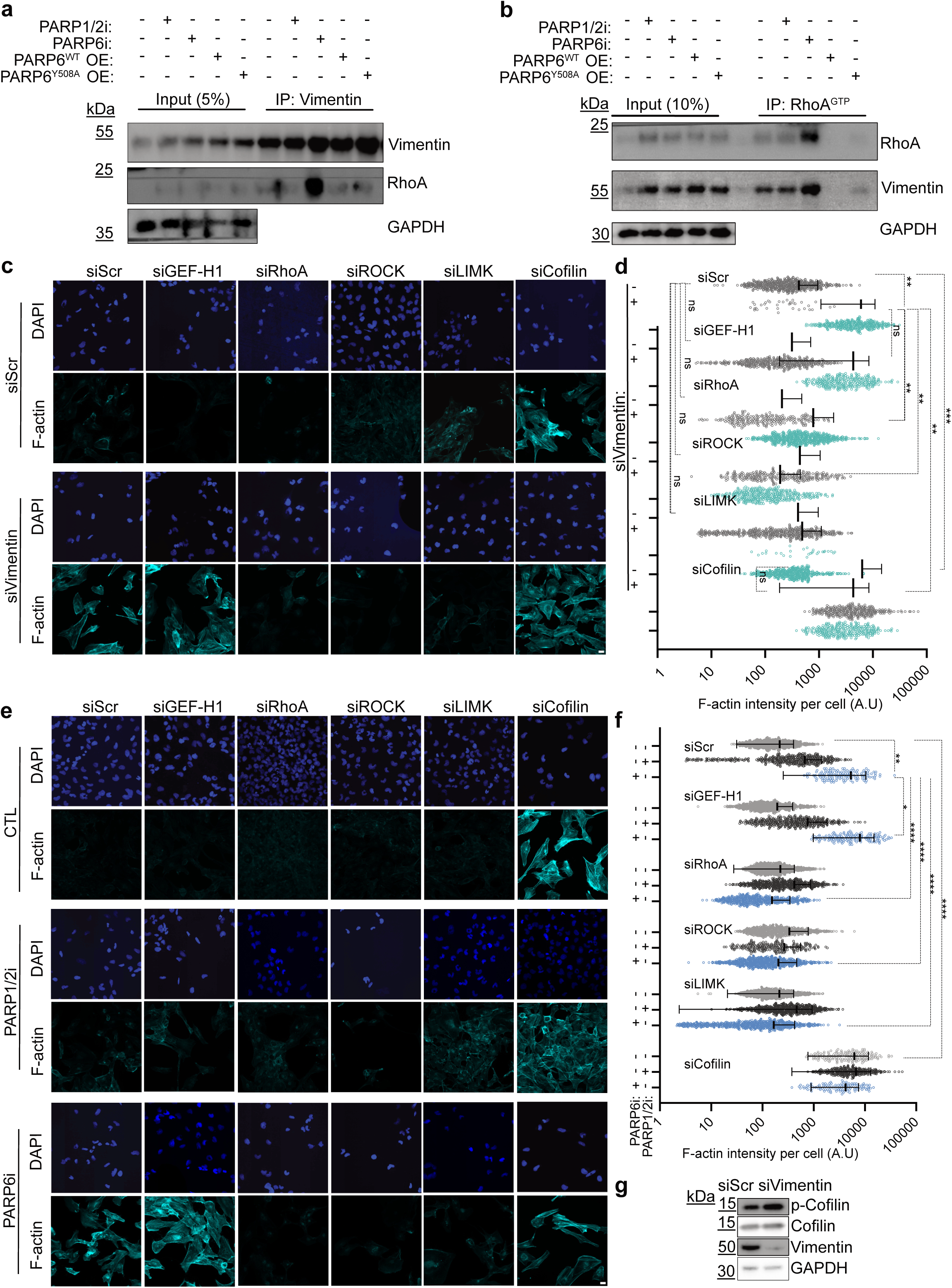
PARP6-dependent vimentin ADP-ribosylation prevents RhoA hyperactivation and restrains RhoA-ROCK-LIMK signaling. **a,** Representative immunoblot of vimentin and RhoA after vimentin immunoprecipitation from cells treated with PARP1/2i (100 nM), PARP6i (1 µM), or overexpression of PARP6^WT^ or PARP6^Y508A^ (induced using 250 ng/ml doxycycline). GAPDH is shown as loading control (n=2). **b,** Representative immunoblot of RhoA and vimentin after rhotekin-based pulldown of RhoA^GTP^ (IP: RhoA^GTP^) from cells treated with PARP1/2i (100 nM), PARP6i (1 µM), or overexpression of PARP6^WT^ or PARP6^Y508A^ (induced by 250 ng/ml doxycycline treatment) as indicated. GAPDH is shown as a loading control (n=2). **c,** Representative confocal microscopy images following knockdown of GEF-H1, RhoA, ROCK1, LIMK, or cofilin under siScr and siVimentin (5nM each) conditions as indicated. Images show DAPI (blue) and F-actin (cyan) (n=3). Scale bar represents 20 µm. **d,** Scatter plot quantification of F-actin intensity per cell corresponding to (**c**), shown for siScr and siVimentin backgrounds. At least 200 cells were counted per condition. Statistical testing was performed on the means of 3 independent experiments using one-way ANOVA with multiple-comparison correction. e, Representative confocal microscopy images following knockdown of GEF-H1, RhoA, ROCK1, LIMK, or cofilin under control, PARP1/2i, or PARP6i treatment as indicated. Images show DAPI (blue) and F-actin (cyan) (n=3). Scale bar represents 20 µm. **f,** Scatter plot quantification of F-actin intensity per cell corresponding to (**e**), shown for siScr and siVimentin backgrounds. At least 200 cells were counted per condition. Statistical testing was performed on the means of 3 independent experiments using one-way ANOVA with multiple-comparison correction. **g,** Representative immunoblot showing P-cofilin, cofilin, and vimentin under siScr and siVimentin treatment conditions. GAPDH is shown as a loading control. (n=2). ns = non-significant, * = p < 0.05, ** = p < 0.01, *** = p < 0.001, **** = p < 0.0001. Scale bar represents 20 µm.

### PARP6 inhibition promotes actin stress fiber organization and myofibroblast activation in primary cardiac fibroblasts

To test whether the PARP6-vimentin-RhoA signaling identified in AC16 cardiomyoblasts is conserved in a disease-relevant cardiac cell type, we tested the mechanism in primary murine cardiac fibroblasts. Vimentin immunoprecipitation from these cells revealed increased co-immunoprecipitation of RhoA with vimentin under PARP6 inhibition, indicating that PARP6 activity similarly constrained vimentin-RhoA complex formation in murine primary cardiac fibroblasts (Fig. 8a) as in human cardiomyoblasts (Fig. 7a). Similarly, immunoprecipitation of RhoA^GTP^ revealed that PARP6 inhibition increased RhoA^GTP^ amounts along with an increase in co-immunoprecipitated vimentin, indicating that vimentin ADP-ribosylation restrained RhoA activation and the preferential interaction of vimentin with RhoA^GTP^ in primary cardiac fibroblasts (Fig. 8b). Consistent with PARP6-mediated regulation of RhoA-ROCK-LIMK-cofilin signaling, PARP6 inhibition increased cofilin phosphorylation while not increasing the total cofilin in primary cardiac fibroblasts (Fig. 8c). This effect of PARP6 pharmacological inhibition was phenocopied by *Parp6*^+/−^ hearts, which showed a marked increase in p-cofilin, along with a moderate increase in the total cofilin expression, further confirming the regulation of RhoA-ROCK-LIMK-Cofilin signaling by PARP6 *in vivo* (Fig. 8d and Fig. S5e).

**Fig. 8:**
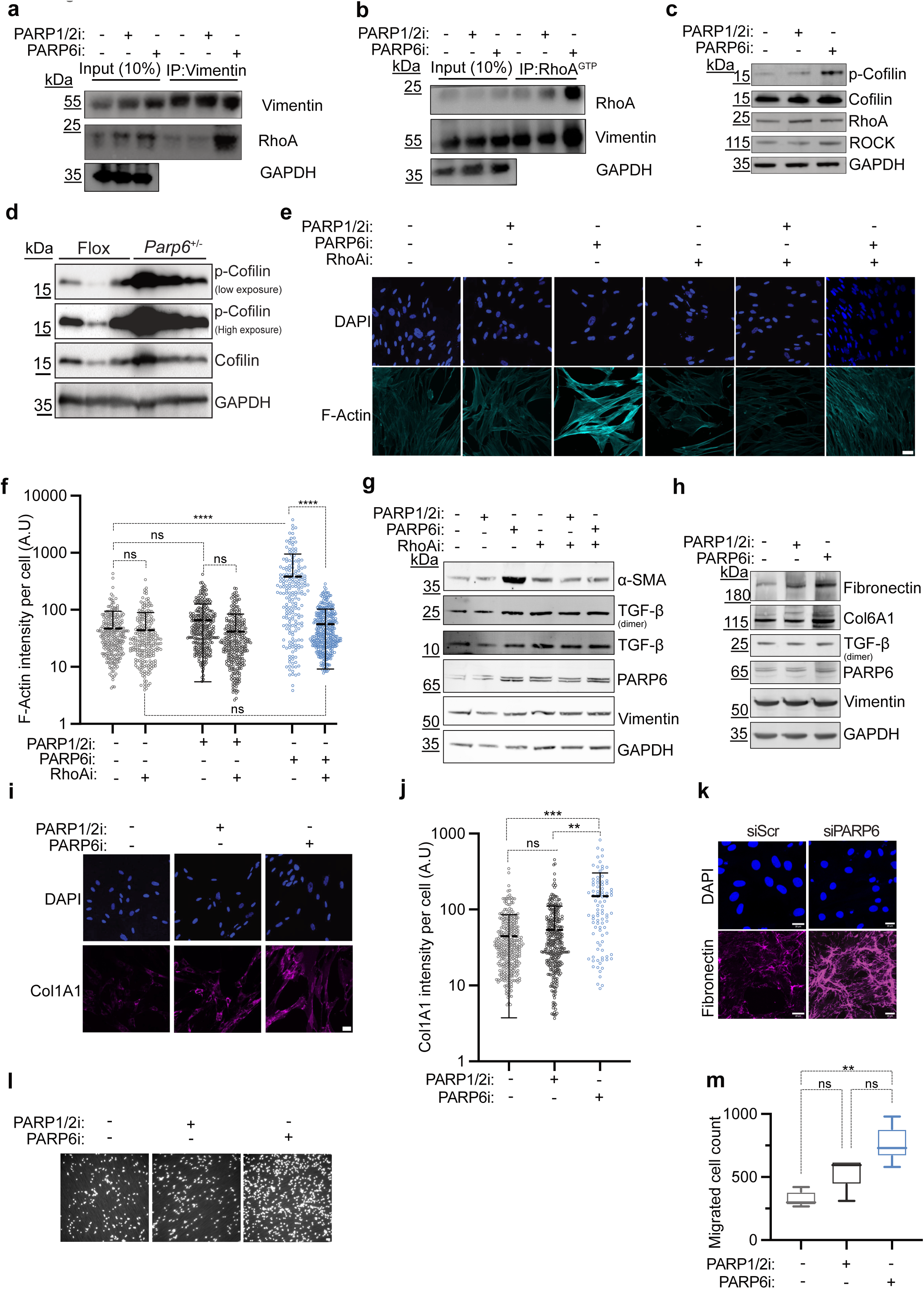
PARP6 inhibition promotes actin stress fiber organization and myofibroblast activation in primary cardiac fibroblasts. **a,** Representative immunoblot of vimentin and RhoA after vimentin immunoprecipitation from primary cardiac fibroblast cells treated with PARP1/2i (100 nM), or PARP6i (1 µM). GAPDH is shown as loading control (n=2). **b,** Representative immunoblot of RhoA and vimentin after rhotekin-based pulldown of RhoA^GTP^ (IP: RhoA^GTP^) from murine primary cardiac fibroblast cells treated with PARP1/2i (100 nM), or PARP6i (1 µM). GAPDH is shown as loading control (n=2). **c,** Immunoblot analysis of murine primary cardiac fibroblasts treated with PARP1/2i (100 nM) or PARP6i (1 µM) as indicated and probed for p-cofilin, total cofilin, RhoA, ROCK, and GAPDH, (n=2). **d,** Immunoblot analysis of p-cofilin and cofilin expression in 6-7 months-old Flox and *Parp6*^+/−^ mice hearts. GAPDH is shown as loading control (n=3). **e,** Representative confocal microscopy images of murine primary cardiac fibroblasts treated with PARP1/2i or PARP6i, with or without RhoA inhibitor (RhoAi) as indicated. Channels: DAPI (blue) and F-actin (cyan), (n=3). Scale bars represent 20 µm. **f,** Violin plot quantification of F-actin intensity per cell from **(e).** At least 200 cells were quantified per condition. Statistical testing was performed on the means of 3 independent experiments using one-way ANOVA with multiple-comparison correction. **g,** Immunoblot analysis of murine primary cardiac fibroblasts treated with PARP1/2i (100 nM), PARP6i (1 µM), or RhoAi (2.5 µM) as indicated and probed for α-SMA, TGF-β, PARP6, vimentin, and GAPDH (n=2). **h,** Immunoblot analysis of murine primary cardiac fibroblasts treated with PARP1/2i or PARP6i as indicated and probed for fibronectin, Col6A1, TGF-β, PARP6, vimentin, and GAPDH (n=2). **i,** Representative confocal microscopy images of murine primary cardiac fibroblasts treated with PARP1/2i (100 nM) or PARP6i (1 µM) as indicated. Images show DAPI (blue) and Col1A1 (magenta) (n=3). Scale bar represents 20 µm. **j,** Violin plot quantification of Col1A1 intensity per cell from **(i)**. At least 75 cells were quantified per condition. Statistical testing was performed on the means of 3 independent experiments using one-way ANOVA with multiple-comparison correction (n=3). **k,** Representative confocal microscopy images of murine primary cardiac fibroblasts following siPARP6 (100 nM), stained for fibronectin (magenta) and DAPI (blue), (n=3). Scale bar represents 10µm. **l,** Representative images from transwell migration assays of murine primary cardiac fibroblasts treated with PARP1/2i (100 nM) or PARP6i (1 µM) as indicated, (n=3). **m,** Bar graph depicting the numbers of migrated cell from **(l)**. Statistical testing was performed on the means of 3 independent experiments using one-way ANOVA with multiple-comparison correction. ns = non-significant. ns = non-significant, * = p < 0.05, ** = p < 0.01, *** = p < 0.001, **** = p < 0.0001.

Having established the conserved signaling, we then tested if there was a PARP6 and RhoA-dependent induction of actin stress fibers in primary cardiac fibroblasts. PARP6 inhibition induced prominent F-actin stress fibers, and the inhibition of RhoA led to a rescue of actin stress fiber induction upon PARP6i treatment, establishing that PARP6i-driven actin remodeling is RhoA-dependent in the cardiac context (Fig. 8e-f).

RhoA activation not only promotes actin stress fiber organization but also integrin-mediated focal adhesion assembly during fibrosis^76^. The cumulative increase in cytoskeletal tension from actin stress fibers and focal adhesions is a key driver of myofibroblast differentiation via mechanosensitive transcriptional programs^77^, such as MRTF–SRF upregulation^78,79^. Functionally, this hypercontractile, actin stress fiber-rich myofibroblast state is linked to increased production of fibrotic ECM components, providing a direct rationale to interpret PARP6-regulated RhoA-driven stress fibers as a proximal step toward fibrosis^89–91^. We therefore tested if the actin remodeling in primary murine cardiac fibroblasts was accompanied by an increase in myofibroblast differentiation markers (Fig. 8g-m). PARP6 inhibition increased α-SMA (a myofibroblast marker)^80^, and the induction of α-SMA was reduced by RhoA inhibition, suggesting that PARP6-regulated cytoskeletal tension promoted myofibroblast differentiation (Fig. 8g). Furthermore, PARP6 inhibition also increased the expression of other ECM remodeling proteins, including fibronectin, collagen6A1 and collagen1A1, suggesting a shift towards a pro-fibrotic, matrix-producing myofibroblast state^67^ restrained by the catalytic activity of PARP6 (Fig. 8h-j). Notably, the increase in myofibrotic ECM proteins was not accompanied by a detectable increase in TGF-β under the same conditions, suggesting that PARP6i promotes fibrotic activation through a TGFβ-independent mechanism, consistent with a contractility- and RhoA-driven activation^79,81^ of myofibrosis (Fig. 8g,h).

To exclude an acute off-target pharmacologic effect of PARP6i, we utilized a targeted siRNA-based approach to deplete PARP6 in primary cardiac fibroblasts. We observed a similar increase in fibronectin (Fig. 8k), confirming that PARP6 restrained myofibrotic transition of cardiac fibroblasts. Since RhoA-dependent cytoskeletal tension is closely coupled to fibroblast motility,^72,82–84^ we assessed the migration of PARP6i-treated cardiac fibroblasts. We observed a significant increase in cellular migration under PARP6 inhibition, while PARP1/2 inhibition resulted in only a moderate increase in migration, further confirming that PARP6 inhibition promoted myofibroblast differentiation, activation, and migration (Fig. 8l-m).

## DISCUSSION

Cardiac fibrosis is increasingly recognized as a maladaptive response that drives heart failure progression^10^, yet the post-translational cues that govern fibroblast activation remain incompletely understood^10^. Our work extends ADP-ribosylation beyond its canonical framing in nuclear stress responses and PARP1-centric PARylation^21,25–27^. Instead, it supports a model in which PARP6-dependent modification functions as a cytoskeletal regulator that couples ADP-ribosylation to mechano-activation in cardiac fibrosis.

A notable feature of the PARP6-dependent ADP-ribosylome is its strong enrichment for cytoskeletal and cytoskeleton-regulatory proteins. While multiple PTMs of vimentin have been described^85–89^, our study, to our knowledge, is the first to identify ADP-ribosylation of vimentin at Cys328 and to implicate PARP6 as its MARylating writer. Importantly, vimentin MARylation altered downstream signaling without detectably perturbing bulk filament architecture, consistent with a model in which ADP-ribosylation rewires vimentin’s interaction landscape rather than its polymerization state. This is conceptually aligned with the emerging view that intermediate filaments function as signaling hubs and that vimentin is not merely a passive scaffold, but a tunable platform that coordinates cross-talk between cellular stress pathways and actomyosin dynamics^45,69,90^.

Cys328 of vimentin is a known hotspot for electrophilic and redox-linked modifications^51^, raising the hypothesis that distinct modifications at Cys328, including MARylation and other electrophilic modifications, could bias vimentin interactomes toward signaling states that either dampen or promote RhoA activation. Because MARylation consumes NAD⁺, this axis also provides a mechanistic link between metabolic state and fibroblast mechanobiology: reduced NAD⁺ availability and PARP6 downregulation may converge to diminish vimentin MARylation, lowering the threshold for RhoA-driven activation during cardiac stress.

Furthermore, PARP6 inhibition induced a coordinated proteomic shift consistent with cytoskeletal remodeling^67^ and fibroblast activation, raising the possibility that this could be linked to PARP6-dependent vimentin MARylation. Loss of vimentin MARylation could favor recruitment of distinct transcription factors/co-regulators to vimentin-associated complexes, reshaping downstream gene-expression programs that reinforce stress-fiber assembly and profibrotic ECM remodeling.

Mechanistically, our data suggest that PARP6 suppresses RhoA activation by preventing the direct interaction of vimentin and RhoA through ADP-ribosylation of vimentin, which has not been described to date. Loss of PARP6 activity increased RhoA^GTP^ levels, enhanced vimentin association with RhoA^GTP^, and activated the RhoA-ROCK-LIMK signaling pathway, leading to phospho-cofilin-dependent actin stress fiber accumulation. The parallel induction of α-SMA and ECM remodeling proteins in primary cardiac fibroblasts, together with phenotypical rescue by RhoA inhibition, indicated that this cytoskeletal shift is sufficient to drive a myofibroblast-like state. A key observation is that PARP6 inhibition in primary cardiac fibroblasts increased α-SMA and extracellular matrix markers without a detectable increase in intracellular TGF-β, whereas TGF-β is elevated in PARP6^+/−^ mouse hearts. This apparent discrepancy suggested that in cell culture models, PARP6 inhibition could trigger a predominantly actin stress fiber-driven, mechanotransductive α-SMA activation program via RhoA-ROCK-LIMK-cofilin signaling, which could precede TGF-β signaling^52,53^. However, in vivo, the myocardium represents a multicellular, complex tissue in which initial cytoskeletal activation and matrix deposition can feed forward into canonical TGF-β-mediated profibrotic signaling^91^. Early ECM accumulation and stiffening can amplify integrin signaling and mechanosensitive transcriptional regulators (e.g., YAP/TAZ- and MRTF-linked programs), and promote activation of latent TGF-β stored in the extracellular matrix through a force-dependent release mechanism^92^. In parallel, additional cell types present in vivo including cardiomyocytes, endothelial cells, infiltrating immune cells, and perivascular populations can provide paracrine sources of TGF-β and other profibrotic mediators in response to injury, inflammation, and hemodynamic stress^93^.

In contrast to the existing literature around PARP1 in cardiac pathology^26^, NAD⁺ consumption by ARTs should not generally be framed as uniformly pathological. PARP1 and tankyrases, which have a very high affinity for NAD^+94^, pose a risk of global NAD^+^ depletion by rapidly generating PAR chains and imposing a high NAD⁺ flux demand during stress. In contrast, PARP6 is a mono-ART^36^, and mono-ADP-ribosylation is intrinsically less NAD⁺ exhaustive per modification site. Moreover, although the affinity of PARP6 for NAD^+^ remains to be defined, ART-family enzymes differ substantially in substrate affinity, catalytic capacity, and abundance supporting a model in which PARP6-mediated MARylation functions as a regulatory layer that promotes cardioprotection.

Our attempts to generate a germline *Parp6* knockout revealed a strong viability constraint. Breeding CMV-Cre-expressing and *Parp6^fl/+^*animals, we generated *Parp6*^+/−^ mice, but repeated reciprocal crosses to *Parp6^fl/fl^* animals failed to yield viable *Parp6*^−/−^ progeny; pregnancies were rare and typically ended in resorption by ∼E15. This aligned with prior evidence that mice expressing a catalytically-inactive, truncated PARP6 variant exhibited severe perinatal lethality or profound postnatal deficits, and that males expressing an inactive/truncated PARP6 were sterile^31,32^. Together with high PARP6 expression in the testis (Biogps.org) and our breeding outcomes, these data support the conclusion that PARP6 is indispensable for development and fertility, underscoring the need for conditional and tissue-specific knockout strategies to interrogate PARP6 function in adult cardiac remodeling.

*In vivo*, *Parp6* haploinsufficiency was sufficient to produce cardiopulmonary and systemic features consistent with pathological remodeling: *Parp6*^+/−^ mice developed cardiac hypertrophy by 6-7 months, as evidenced by increased heart and lung weights. The increased lung weight was consistent with reports linking cardiac dysfunction to lung remodeling, fibrosis, and type 2 pulmonary hypertension^95^. We also observed splenomegaly, consistent with bidirectional spleen-heart axis regulation, in which splenic immune activation can both reflect and amplify chronic inflammation, tissue injury, and cardiac remodeling^96^. Therapeutically, the convergence on RhoA-ROCK-LIMK highlighted mechano-transduction inhibition as a rational anti-fibrotic strategy and suggested caution regarding systemic PARP6 inhibition, which could inadvertently bias fibroblasts toward activated myofibroblast phenotypes. Therapeutically, the convergence on RhoA-ROCK-LIMK highlighted mechano-transduction inhibition as a rational anti-fibrotic strategy and suggested caution regarding systemic PARP6 inhibition, which could inadvertently bias fibroblasts toward activated myofibroblast phenotypes.

### Limitations of the study

The upstream GAP/GEF proteins responsible for RhoA activation in a vimentin- ADP-ribosylation-dependent manner remain undefined. An intriguing possibility remains that PARP6 inhibition may not reflect the changes in the interaction of a single GAP/GEF with RhoA in a vimentin ADP-ribosylation-dependent manner, but rather a shift in the stoichiometry and competitive complexing of multiple GEFs and GAPs with RhoA. This could alter the balance of activation versus inactivation without requiring a single RhoA regulator to be uniquely responsible for PARP6-induced myofibroblast transition. Future interactome studies may identify other RhoA regulators or a MARylation-dependent effector (reader) that links vimentin ADP-ribosylation to RhoA regulation.

## MATERIALS AND METHODS

### Human heart samples

All research related to human patients and samples was reviewed and approved by the Institute’s human ethical committee (IHEC approval No: 20/5.8.2021), along with the ethical committee of Sri Jayadeva Institute of Cardiovascular Sciences and Research (IHEC approval no. SJICR/EC/2021/039), and the use of human biological material in Switzerland was approved by the cantonal ethics committee of the Canton of Zurich (BASEC-Nr. 2021-02162). Cardiac functional parameters were evaluated in patients enrolled in the hospital’s surgical unit, according to standard AHA/ACC, European, and Indian guidelines. Both males and females were recruited to the study. These patients were subsequently grouped into different categories based on the clinical manifestations of heart failure. Patients were identified based on the degree of heart failure, with patients having left heart failure with associated right ventricular failure being enrolled. All patient identities were anonymized, and the team evaluating tissue histology and molecular biology was blinded to the nature of the reasons for heart failure. After certification from the treating cardiologist for the need for surgery, informed consent was obtained from patients, and they were placed on a conventional cardiopulmonary bypass. An endomyocardial biopsy, with an endomyocardial bioptome/punch taken from the LV endomyocardium of these patients. Control samples were collected as biopsies from hearts that were donated for homograft banking, or from forensic autopsies of individuals with no history or pathological evidence of pre-existing heart failure, i.e., death due to unrelated causes. The heart tissue samples were snap-frozen in liquid nitrogen and stored at −80°C. Samples were also collected in 10% neutral buffered formalin for histological analysis. Detailed information on the cardiac parameters of patients is presented in Table 1.

**Table 1:**
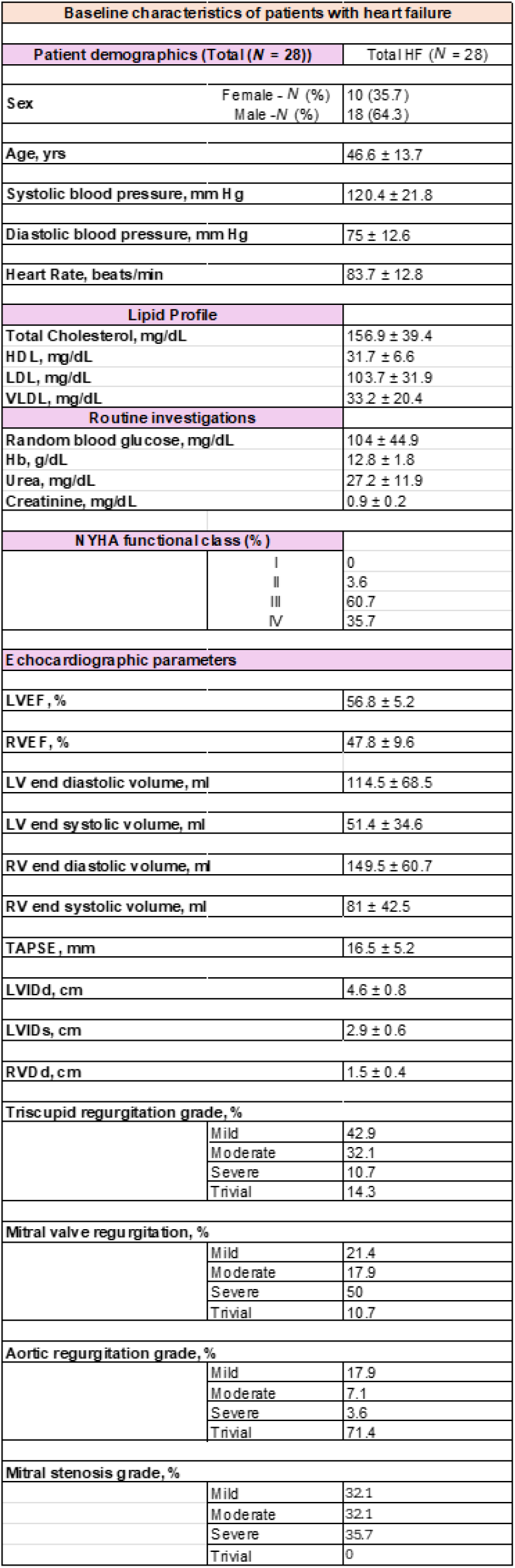
Values are mean ± SD unless stated otherwise. HF = Heart failure. HDL= High density lipoprotein. LDL= Low density lipoprotein. VLDL= Very low-density lipoprotein. Hb= Hemoglobin. NYHA=New York Heart Association. LVEF = Left ventricular ejection fraction. RVEF = Right ventricular ejection fraction. TAPSE = Tricuspid Annular Plane Systolic Excursion. LVIDd= Left Ventricular Internal Diameter at End-Diastole. LVIDs= Left Ventricular Internal Diameter at End-systole. RVDd= Right Ventricular End-Diastolic diameter.

### Animal studies

Animal handling and associated experiments were performed after obtaining due approval by the Institutional Animal Ethics Committee (IAEC) of the Indian Institute of Science, in compliance with guidelines of article 13 of the Committee for the Purpose of Control and Supervision of Experiments on Animals (CPCSEA), Government of India. All studies performed were verified for compliance with the Guide for the Care and Use of Laboratory Animals, US National Institute of Health (NIH Publication No. 85–23, revised 1996). Animals were housed in a clean air, pathogen-free facility and fed a standard chow diet, ad libitum food and water, with a 12h light/dark cycle at the Central Animal Facility, Indian Institute of Science. *Parp6^fl/fl^* and *CMV-Cre* mice used were procured from Cyagen US Inc. Global *Parp6^+/−^* were generated by crossing male *PARP6^fl/fl^* mice with female *CMV-Cre* mice to obtain *Parp6^fl/fl^ CMV-Cre*^+^ (or *Parp6^+/−^*) mice. Upon generation, we attempted to backcross these *Parp6^+/−^* animals to *Parp6^fl/fl^*mice to generate knockout (*Parp6^−/−^*) mice. Intriguingly, despite reciprocal crossing, we were unable to obtain any viable progeny from this breeding. Specifically, the breeding of *Parp6^+/−^* male with *Parp6^fl/fl^* female did not result in a pregnancy. In crosses involving female *Parp6^+/−^* mice and *Parp6^fl/fl^* male, pregnancy was only rarely observed and typically resulted in resorption by approximately day 15 of gestation, consistent with recent findings^32^

All *in vivo* studies on *Parp6^fl/fl^*, *CMV-Cre* and *Parp6^+/−^* mice were performed on 6-7-month-old male mice unless stated otherwise. Age-matched male *Parp6^fl/fl^* mice were used as controls. Primary investigations were performed with male mice (unless mentioned for female mice data), as males exhibit increased susceptibility to cardiac dysfunction, including increased vulnerability to heart failure and hypertrophic remodeling. Conversely, females experience cardioprotection due to the pleiotropic effects of estrogen, including reduced cardiac ROS production, improved calcium handling and altered inflammatory signaling^97^. Mice tibia lengths were measured using vernier caliper. The heart, liver, kidney, spleen and brain were harvested from mice, weighed and snap-frozen in liquid nitrogen.

For the development of a murine model of experimentally induced hypertrophy, 3 to 4-month-old C57BL/6 male mice were injected intraperitoneally either with vehicle or ISO (10 mg/kg of body weight) daily for 14 days. Cardiac function was assessed by echocardiography using the Vevo® 1100 Imaging System (FUJIFILM VisualSonics).

### Echocardiography of mice

Mice were anesthetized with continuous ∼1% isoflurane delivered via a nasal cone. Chest hair was removed using a commercial topical depilatory, and body temperature was maintained with a heated imaging platform. Electrocardiogram (ECG) leads were attached to the limbs for cardiac gating. A FUJIFILM VisualSonic Vevo 1100 high-frequency ultrasound system with a 30 MHz transducer was used to image the mice in the left lateral decubitus position. Two-dimensional echocardiographic images were recorded in parasternal long- and short-axis views, with M-mode recordings guided at the midventricular level in both views. Left ventricular cavity size and wall thickness were measured from at least three cardiac cycles in each view and averaged for analysis. Left ventricular ejection fraction and fractional shortening were calculated from M-mode measurements using the system’s built-in software.

### Cardiac histology

Age-matched *Parp6*^fl/fl^ and *Parp6*^+/−^ hearts were collected and fixed in 10% neutral buffered formalin. After fixation, hearts were embedded in paraffin wax and then cut into sections of 5 µm thickness. An automated tissue processer (Leica) was used for tissue processing. Fibrosis was observed and measured by staining paraffin-embedded sections with Masson’s trichrome stain (Solarbio, G1340). Cardiomyocyte size was measured in sections stained with Wheat Germ Agglutinin (WGA; 5 μg/ml). Masson’s trichrome staining was performed for control and failing human samples using the same procedure.

### Isolation of neonatal rat cardiac fibroblasts

Primary culture for neonatal rat cardiac fibroblasts was performed as described previously^98^. Briefly, hearts were harvested from 0-2 days old neonatal rat pups and collected in a solution of 1X phosphate buffer saline supplemented with glucose (PBSG). Tissues were minced with a sterile surgical blade and digested in a mixture containing 0.25% trypsin and 0.4 mg/mL collagenase II by constant rotation for 5-10 minutes at 37°C in a shaker incubator. After every digestion cycle, the supernatant containing the digested cells was collected in horse serum. This process was repeated for 7-10 cycles. The collected cell mixture was then pre-plated on a 100mm dish for 1 hour at 37°C, after which, the supernatant containing cardiomyocytes was separated and only the attached cells were taken for further experiments. The cells were grown and maintained in high-glucose DMEM (Dulbecco’s Modified Eagle Medium) supplemented with 10% Fetal Bovine Serum (FBS) and 1X antibiotic-antimycotic cocktail at 37°C and 5% CO_2_. These cells were then trypsinized and seeded into appropriate gelatin-coated dishes for the experiments mentioned.

### Isolation of murine primary cardiac fibroblasts

Primary murine cardiac fibroblasts were isolated with minor adaptations from a protocol previously described^99^. Briefly, murine hearts were excised, rinsed in cold phosphate-buffered saline (PBS), and ventricles were minced into small fragments, which were digested in collagenase/DNase-based digestion cocktail (Liberase Blendzyme 3 or Liberase TH (Roche) supplemented with DNase I and HEPES in HBSS containing Ca²⁺/Mg²⁺). The digested suspension was diluted in PBS and passed through a 40 µm strainer to obtain a single-cell suspension. Cells were pelleted, subjected to red blood cell lysis for 5 minutes, washed, re-filtered, and plated in high-glucose DMEM containing 10% FCS, 1% penicillin/streptomycin, and 20 µg/ml plasmocin (Invivogen). Cells were incubated at 37 °C to allow selective adhesion, and after 4 hours, non-adherent cells were removed. Adhered fibroblasts were expanded and used for further experiments.

### Cell culture

AC16 cardiomyoblasts were obtained from ATCC (CRL-3568). Wild-type and vimentin^−/−^ mouse MEF cells were a kind gift from Prof. Ohad Medalia. Primary cardiac fibroblasts were isolated from mice. AC16 cells were maintained in phenol red free Dulbecco’s modified Eagle’s medium/ F12 (DMEM/F12) supplemented with 12.5% fetal calf serum and 1% penicillin/streptomycin. MEFs and primary cardiac fibroblasts were cultured in high-glucose Dulbecco’s modified Eagle’s medium (DMEM) supplemented with 10% fetal calf serum (FCS) and 1% penicillin/streptomycin. All cell types were grown at 37 °C in a humidified incubator with 5% CO₂ and were routinely tested for mycoplasma contamination. Cells were treated for 24 hours with either talazoparib (100nM) or AZ0108 (1µM; a kind gift from AstraZeneca), PJ34 (10 µM), or bafilomycin A1 (100 nM) unless indicated otherwise. For experiments using ROCK and RhoA inhibitors, cells were treated with either Y27632 (ROCKi, 10 µM) for 1 hour or Rhosin Hydrochloride (RhoA inhibitor, 2.5 µM) for 24 hours. For vimentin filament formation analysis, cells were serum starved overnight and treated with diamide (100 µM) for 1 hour, or talazoparib (100 nM), AZ0108 (1 µM), or doxycycline (250 ng/mL) to induce PARP6 overexpression for 24 hours.

### PARP6 overexpression in AC16 cells

Stable genomic integration of PARP6 in AC16 cardiomyoblasts was performed using the Sleeping Beauty transposon system. Briefly, AC16 cells were co-transfected with a Sleeping Beauty donor vector (Addgene plasmid #60506) cloned to encode the coding sequence of PARP6 and the Sleeping Beauty transposase plasmid (Addgene plasmid #34879) using Lipofectamine 3000 (Thermo Fisher Scientific). At 72 hours post-transfection, cells were placed under 1μg/mL blasticidin selection to obtain stably expressing populations. PARP6 overexpression was induced using 250ng/mL doxycycline for 16 hours.

### siRNA mediated knockdown

siRNA-mediated gene silencing was carried out using Lipofectamine RNAiMAX (Thermo Fisher Scientific). Briefly, siRNAs (synthesized by Microsynth AG) were diluted to a final concentration of 10 nM or 100 nM as indicated in serum-free Opti-MEM and combined with Lipofectamine RNAiMAX (5 µL per well of a 6-well plate) in a total volume of 500 µL, followed by incubation at room temperature for 15 minutes. The transfection complexes were then added dropwise to freshly adhered cells seeded in 6-well plates, and the medium was replaced 12 hours post-transfection. A non-targeting scrambled siRNA (siScr) was included in all experiments as a negative control. For co-depletion experiments, the combined siRNA concentration was maintained at 10 nM. The siRNA sequences (5’-3’) used include siScr: UUC UCC GAA CGU GUC ACG UUU, siVimentin: UCA CGA UGA CCU UGA AUA ATT, siGEFH1: GGA CAA GCC UUC AGU GGU ATT and GAA UUA AGA UGG AGU UGC ATT, siRHOA: UUC UGU ACG AAC GAG UAU CAG and GAC AUG CUU GCU CAU AGU CUU, siROCK: CAC CGC GGA GGA AGU UGG UUG AAA UTT, GCA GCA AUG GUA AGC GUA ATT, GCA ACU GGC UCG UUC AAU UTT, and UGG UGC UGG UAA GAG GGC AUU GUC ATT, siLIMK: GCA AGC GUG GAC UUU CAG UTT and AAA CCG GAU CUU GGA AAU CTT, siCofilin: CCA CCU UUG UCA AGA UGC UTT and GAU CAA GCA UGA AUU GCA ATT, siPARP6: GGC GAU GCC AAC AUU AAU ATT, UAU UAA UGU UGG CAU CGC CTT, and Thermo Silencer siPARP6 (AM16708).

### Immunoprecipitation of vimentin

Cells were treated with DMSO, PARP1/2i (100nM talazoparib), PARP6i (1µM AZ0108) for 24 hours. Following treatment, cells were lysed using Pierce lysis buffer (25 mM Tris-HCl (pH 7.4), 150 mM NaCl, 1% NP-40, 1 mM EDTA, 5% glycerol, and protease inhibitors) and incubated at 4 °C under constant rotation for 20 min. The lysates were then clarified by centrifugation at 14,000 × g for 10 min at 4 °C. Dynabeads Protein G (Thermo Fisher Scientific) were incubated with anti-vimentin (clone V9; Santa Cruz Biotechnology, sc-6260), or mouse IgG control antibody (Millipore) for 2 hours at 4 °C with rotation for reactions from AC16 cardiomyoblasts. Beads were then washed and equilibrated in wash buffer (25 mM Tris-HCl, pH 7.4, 150 mM NaCl, 1 mM EDTA, and protease inhibitors) and then combined with clarified cell lysates for 1 hour at 4 °C with rotation. Following incubation, beads were washed three times with wash buffer. For ADP-ribosylation of vimentin, the final wash included 5% 1,6-hexanediol (prepared in wash buffer) to increase stringency before elution. Beads were then resuspended in 30 µL SDS sample buffer, boiled, and immunoblotted.

### Immunoprecipitation of RhoA^GTP^

RhoA^GTP^ was isolated using a rhotekin-Rho binding domain (RBD) affinity pull-down (Rho Activation Assay Biochem Kit, Cytoskeleton Inc., BK036). Following the indicated treatments, cells were rapidly transferred to ice, washed with ice-cold PBS, and lysed in ice-cold kit Cell Lysis Buffer supplemented with protease inhibitors. Lysates were clarified by centrifugation (10,000 × g, 1 min, 4 °C), and protein concentrations were measured and equalized across conditions. 800 µg total protein was incubated with 50 µg rhotekin-RBD for 1 h at 4 °C with rotation. Beads were pelleted (5,000 × g, 1 min, 4 °C), washed once with kit Wash Buffer, and re-pelleted (5,000 × g, 3 min, 4 °C). Bound proteins were eluted by resuspending beads in 2× Laemmli sample buffer, boiling for 10 min, and analyzed by immunoblotting.

### Immunoblotting

Cells were lysed in SDS/LDS lysis buffer (60 mM Tris-HCl, pH 7.1, 2% SDS/LDS, and 10% glycerol). Lysates were denatured at 70 °C for 10 min and briefly sonicated to reduce viscosity. Proteins were resolved on 12% SDS-PAGE or Bis-Tris polyacrylamide gels (120 V) and transferred to PVDF membranes by wet transfer (100 V, 2 h). Membranes were blocked in 5% (w/v) milk/TBS-T for 1 hour at room temperature and incubated with primary antibodies diluted in 5% milk/TBS-T overnight at 4 °C. After washing four times with TBS-T, membranes were incubated with appropriate infrared secondary antibodies diluted in 5% milk/TBS-T for 1 h at room temperature, washed an additional four times, and imaged.

Primary antibodies used for immunoblotting included: vimentin (clone V9, Santa Cruz Biotechnology, sc-6260, 1:2000; Cell Signaling Technology, #5741, 1:2000), PARP6 (Thermo Fisher Scientific, #31988, 1:2000), GAPDH (Cell Signaling Technology, #2118, 1:5000), RhoA (Cell Signaling Technology, #2117, 1:1000), ROCK1 (Cell Signaling Technology, #28999, 1:1000), cofilin (Cell Signaling Technology, #5175, 1:1000), phospho-cofilin (Ser3) (Cell Signaling Technology, #3313, 1:1000), α-smooth muscle actin (Thermo Fisher Scientific/eBioscience, #14-9760-82, 1:100), TGF-β (Cell Signaling Technology, #3711, 1:1000), fibronectin (Cell Signaling Technology, #26836, 1:1000), and collagen VIα1 (Cell Signaling Technology, #58232, 1:1000). Western blot images were acquired in Amersham Typhoon or Witec Fusion 2 imagers, or using X-Ray films.

For *in vivo* samples, tissues were lysed in ice-cold tissue lysis buffer (20 mM Tris-HCl, pH 7.5, 150 mM NaCl, 1 mM EGTA, 1 mM EDTA, 1% Triton X-100, 1% Sodium deoxycholate, 2.5 mM sodium pyrophosphate, 1 mM sodium orthovanadate, 1 mM PMSF, 1mM Sodium Fluoride, and 1X protease inhibitor cocktail). Lysates were cleared by centrifugation at 13,000 rpm for 15 min at 4°C and the supernatant was collected in a fresh tube.

### Transwell migration assay

Cell migration was assessed using transwell assay with collagen coated polycarbonate membrane inserts (8.0 µm pore size; Corning). Primary cardiac fibroblasts were seeded into the upper chamber and treated with either a PARP1/2 inhibitor (100 nM) or a PARP6 inhibitor (1 µM). Cells were allowed to migrate for 48 hours under standard culture conditions. Following incubation, non-migrated cells remaining on the upper surface of the membrane were gently removed using a cotton swab. Migrated cells on the lower surface of the membrane were stained with DAPI (0.1 µg/mL in PBS) for 20 min at room temperature. Membranes were washed with PBS and imaged using a THUNDER microscope (Leica). Migrated cells were quantified by counting DAPI-positive nuclei on the lower surface of the membrane.

### Quantitative Real Time PCR

Total RNA was extracted from cells using RNAiso Plus (Takara Bio) and subsequently purified by a phenol–isopropanol–ethanol method. After verification of RNA concentration and integrity, 1 μg of RNA was reverse transcribed in a 20 µl reaction volume to generate cDNA using the PrimeScript™ 1st Strand cDNA Synthesis Kit (Takara Bio). 1:50 and 1:100 dilutions of the resulting cDNA (3-4 µl per reaction) were then employed in real-time PCR to evaluate the expression of selected genes. RT-PCR was carried out using TB Green® Master Mix (Takara Bio), and amplification was performed on a QuantStudio 6 Flex System (Thermo Fisher Scientific). Human primer sequences hPARP3(Forward): GGACTGTGACTAAGCGGGTG, hPARP3(Reverse): TGAGGGCCATGGTGTTCTTG; hPARP10(Forward): TTGAAGAGGTGGACCCTACCG, hPARP10(Reverse): CAGTCCTCCCCGTCTAGGTC; hPARP8 (Forward): GCAGGTCGCCAAGTTATCCT, hPARP8 (Reverse): AGGGCAAGGACGGTTCAAAA; hPARP9 (Forward): GACCCTACTGTTGCTGCCTT, hPARP9 (Reverse): ATGTGGCCCTGGACAATCTG, hPARP6(Forward): CCATGCTGAAAGGCAAACTAGG, hPARP6(Reverse): TGGTACAGGAGATGGAACGGT; hGAPDH(Forward): ACAACTTTGGTATCGTGGAAGG, hGAPDH (Reverse): GCCATCACGCCACAGTTTC. Murine primer sequences: mPARP6(Forward): CGATGACGTGGACCTTGACC, mPARP6 (Reverse): TCGATGGCTGGAAAACCTCAA; mGAPDH(Forward): TATGTCGTGGAGTCTACTGGT, mGAPDH (Reverse): GAGTTGTCATATTTCTCGTGG

### Immunofluorescence microscopy

For immunofluorescence (IF), cells were seeded on coverslips and subjected to the indicated treatments. Cells were fixed with 4% formaldehyde in PBS for 20 min at room temperature and permeabilized in PBS containing 0.2% Triton X-100 (Sigma-Aldrich) for 15 min at room temperature. Following blocking in PBS supplemented with 0.1% Triton X-100 and 2% bovine serum albumin (BSA) for 2 hours, coverslips were incubated with primary antibodies diluted in blocking buffer overnight at 4 °C. Coverslips were washed three times with PBS and incubated with fluorophore-conjugated secondary antibodies diluted in blocking buffer for 1 h at room temperature. Nuclei were counterstained with DAPI (0.1 µg/mL in PBS) for 20 min at room temperature. Coverslips were washed three times in PBS (5 min each), mounted on glass slides using Mowiol, and imaged on a confocal microscope.

Primary antibodies/reagents used for IF included: vimentin (clone V9, Santa Cruz Biotechnology, sc-6260, 1:250; Cell Signaling Technology, #5741, 1:250), phalloidin-Alexa Fluor 488 (Abcam, ab176753, 1:800), LAMP1 (BioLegend, #328602, 1:250), anti-HA (Cell Signaling Technology, #2367, 1:250), and collagen Iα1 (Cell Signaling Technology, #72026, 1:250), Fibronectin (Santa Cruz Biotech C-20, #6952, 1:50), Secondary antibodies used were goat anti-rabbit IgG (H+L) Alexa Fluor 647 (Invitrogen, A21244, 1:500) and goat anti-mouse IgG (H+L) Alexa Fluor 488 (Invitrogen, A11029, 1:500). Leica thunder, Olympus spinning disc, Zeiss Airyscan, Leica SP8 Falcon and Andor Dragonfly Spinning Disk Confocal Microscopes were used to acquire images.

For super-resolution imaging, cells were seeded on high-precision #1.5H coverslips (170 ± 5 µm thickness) compatible with high-NA oil-immersion objectives. Following staining and final PBS washes, coverslips were mounted onto glass slides using ProLong Diamond Antifade Mountant (Invitrogen, P36961) and protected from light prior to acquisition in Zeiss Elyra 7.

### Image intensity and radial distribution analysis

Fluorescence image analysis was performed using CellProfiler (v4.1, BROAD Institute). Images were first corrected for uneven illumination and background using the CorrectIllumination and SubtractBackground modules. Individual cells were identified based on the nuclear channel using the IdentifyPrimaryObjects module, followed by the identification of whole-cell boundaries using the IdentifySecondaryObjects module. Mean fluorescence intensity was quantified for each cell using the MeasureObjectIntensity module. To assess the radial distribution of fluorescence, cell objects were divided into concentric radial bins extending from the center of the nucleus to the cell periphery using the MeasureObjectRadialDistribution module. The coefficient of variation (CV) of radial intensity was calculated for each cell as the standard deviation of fluorescence intensity across radial bins divided by the mean intensity. Single-cell measurements were exported and aggregated for statistical analysis.

### Immunofluorescence of neonatal rat primary cardiac fibroblasts

Cells were fixed with formaldehyde (4%) and were permeabilized using 0.2% Triton X-100. Cells were incubated with the primary antibodies, followed by incubation with secondary antibodies conjugated with Alexa Fluor 488 and/or 546. Hoechst was used to stain the nuclei. An Andor Dragonfly Spinning Disk Confocal Microscope was used to acquire images.

### Cell lysates for mass spectrometry

Cells were lysed in denaturing lysis buffer (6 M guanidine hydrochloride, 50 mM Tris-HCl, pH 8.0), sonicated, snap frozen, and stored at −80 °C until LC-MS/MS analysis. Disulfide bonds were reduced with 10 mM tris(2-carboxyethyl)phosphine (TCEP) and alkylated with 20 mM 2-chloroacetamide (CAA) for 30 min at 30 °C in the dark. For whole-proteome analysis, 50 µg of total protein was processed using filter-aided sample preparation (FASP) and digested with sequencing-grade trypsin (enzyme:substrate 1:25; Promega) overnight at 37 °C. Digests were acidified with trifluoroacetic acid (TFA) and desalted using C18 ZipTip pipette tips (Millipore). Peptides were eluted in 15 µl of 60% acetonitrile (ACN)/0.1% TFA, dried to completion, and reconstituted in 3% ACN/0.1% formic acid to a final concentration of 0.5 µg/µl. For ADP-ribosylome analysis, 10 mg of protein was diluted 1:12 in PARG buffer (50 mM Tris-HCl, pH 8, 50 mM NaCl, 10 mM MgCl2, 250 µM DTT) and digested with modified porcine trypsin (enzyme:substrate 1:25; Sigma) overnight at 37 °C. ADP-ribosylated peptides were enriched as described previously^37^, with the following modifications: after PARG-mediated conversion of PAR to MAR, peptides were incubated with Af1521^WT^ (0.5 ml beads per 15 mg lysate) and eAf1521 (1.0 ml beads per 15 mg lysate) for 2 h at 4 °C. Enriched fractions were subsequently processed for MS analysis as described ^37^.

### Liquid chromatography-tandem mass spectrometry (LC-MS/MS)

ADP-ribosylated peptides were analyzed on an Orbitrap Lumos mass spectrometer (Thermo Fisher Scientific) coupled to an ACQUITY M-class UPLC (Waters) using an ADP-ribose product-dependent method (HCD-PP-EThcD). Mobile phases were 0.1% formic acid in water (A) and 0.1% formic acid in acetonitrile (B). Peptides were loaded onto a nanoEase M/Z Symmetry C18 trap column (18 µm × 20 mm, 5 µm, 100 Å; Waters) and separated on a nanoEase M/Z HSS T3 analytical column (75 µm × 200 mm, 1.8 µm, 100 Å; Waters) at 300 nl/min over 110 min using the following gradient: 3–25% B for 95 min, 25–35% B for 5 min, and a wash at 95% B for 5 min. Full-scan MS1 spectra (350–2000 m/z) were acquired at 120,000 resolution (AGC target 4 × 10^5^; maximum injection time 50 ms). HCD MS/MS spectra (collision energy 38%) were acquired at 30,000 resolution (AGC target 5 × 10^4^; maximum injection time 60 ms). Detection of at least two diagnostic ADP-ribose fragment ions (136.0623, 250.0940, 348.0709, and 428.0372 m/z) in the HCD scan triggered additional high-resolution HCD and EThcD MS/MS scans (120,000 resolution; AGC target 5 × 10^5^; maximum injection time 240 ms; HCD collision energy 35%).

MS data analysis: Raw MS1 and MS2 spectra were converted to Mascot generic format (MGF). Separate MGF files were generated for HCD and EThcD spectra. Mascot searches were performed as described previously against the UniProtKB human database (taxonomy 9606; version 20190709), including Swiss-Prot and TrEMBL entries, decoys, and common contaminants. Carbamidomethylation of cysteine was set as a fixed modification, while protein N-terminal acetylation and methionine oxidation were specified as variable modifications. ADP-ribosylation was included as a variable modification on C, D, E, H, K, R, S, T, and Y residues. For HCD spectra, neutral losses of 347.0631 Da and 583.0829 Da were included for scoring of ADP-ribose-derived fragment ions.

## DATA AVAILABILITY

Original data is available from the authors upon request. The mass spectrometry proteomics data have been deposited to the ProteomeXchange Consortium via the MassIVE partner repository with the dataset identifier PXD074340. Download link via MassIVE: ftp://massive-ftp.ucsd.edu/v12/MSV000100800/.

## ACKNOWLEDGMENTS

We thank Flurina Böhi, Patrick Manetsch, and Tobias Suter (University of Zurich) for valuable discussions and editorial support. We also thank John Eriksson (University of Turku) for scientific input. Vimentin^−/−^ MEF cells were a gift from Prof. Ohad Medalia (University of Zurich). We thank the Center for Microscopy and Image Analysis (ZMB) and the Functional Genomics Center Zurich (FGCZ), University of Zurich, for their technical services and assistance. We further thank Peter Gehrig and Jonas Grossmann (FGCZ) for insightful discussions, methodological recommendations, and bioinformatics support. We also sincerely thank Dr. Jamie Scott (AstraZeneca) for providing AZ0108. Fig. 3a was created in BioRender: Sundaresan, S. (2026) https://BioRender.com/6gbnevk. Fig. 4a was created in BioRender: Sundaresan, S. (2026) https://BioRender.com/ftghtx9. Supplementary Fig. 2a was Created in BioRender: Sundaresan, S. (2026) https://BioRender.com/3kdaigx. This study was executed under the Indo-Swiss Joint Research Program financed by a grant from the Swiss National Science Foundation to MOH (grant IZLIZ3_200237) and from the Department of Biotechnology (IC-12044[11]/3/2021-ICD-DBT) to NRS.

## AUTHOR CONTRIBUTIONS

Funding Acquisition: MOH, NRS; Conceptualization: SS, AT, MOH, NRS; Data curation: SS (cell culture models), AT(animal models); Formal analysis: SS (cell culture models), AT (animal models); Methodology: SS (cell culture models), AT (animal models), DK (microscopy), DMLP (mass spectrometry and analysis); Project administration MOH, NRS; Software: SS, AT; Supervision: MOH, NRS; Writing original draft preparation: SS (cell culture models), AT (animal models); Review and Editing: MOH, NRS, SS, AT.

---

**Supplementary Fig. 1:**
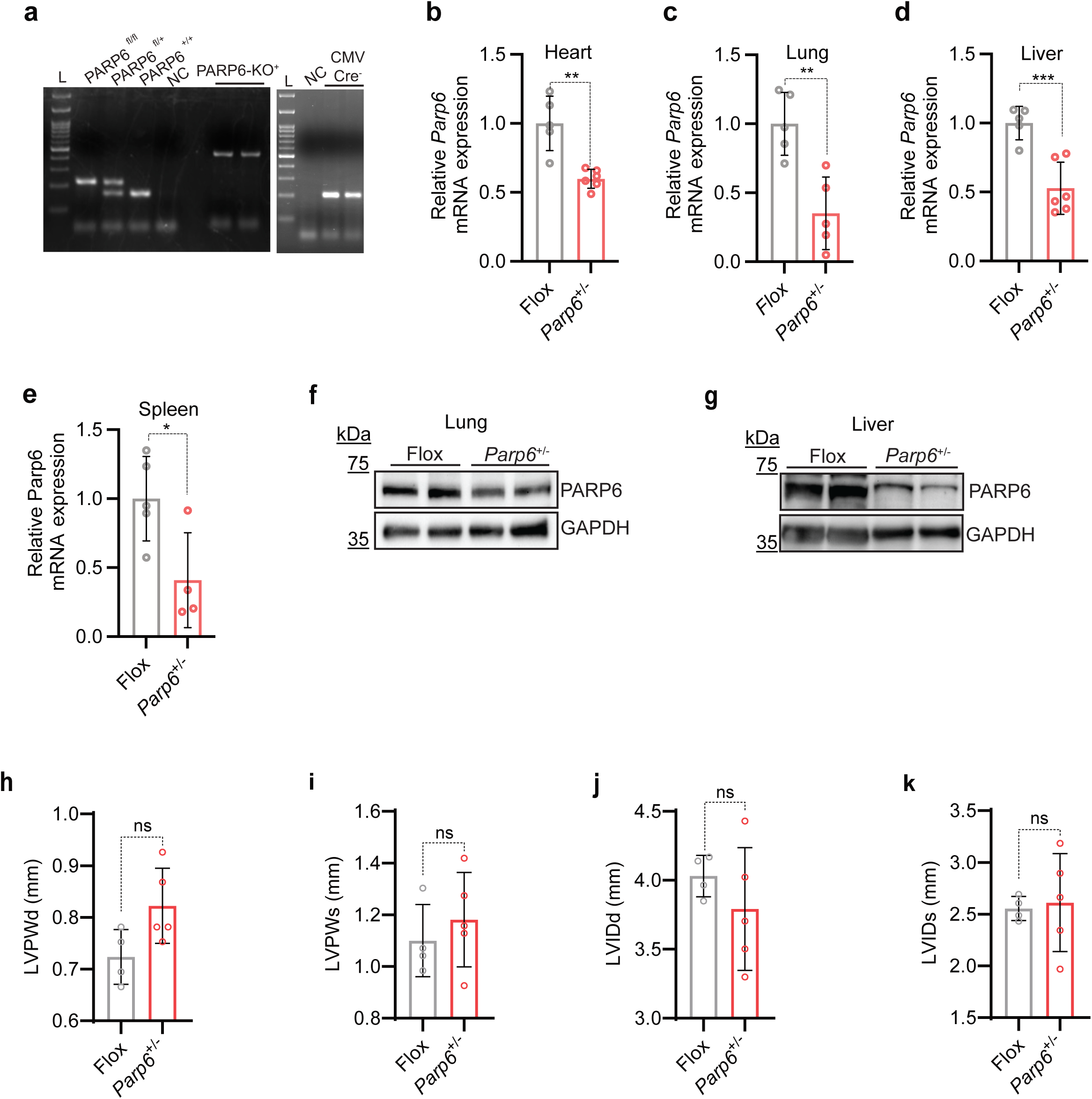
Whole-body PARP6^+/−^ mice generation and validation. **a,** Representative image of genomic PCR-based confirmation of PARP6 global deletion by screening for a *Parp6*^fl/+^ CMV-Cre^+^ combination of mice. The band at 226 bp represents the flox band, while the band at 166 bp represents the wild-type band. Heterozygous *Parp6*^fl/+^ mice exhibit amplification of both bands. Mice positive for CMV-Cre show amplification at 204 bp. CMV-Cre-positive *Parp6*^fl/+^ mice show amplification at 439 bp in the *Parp6*-KO primer set (*Parp6*-KO^+^), representing the *Parp6* allel with deleted exons 5-8 in the whole body (*Parp6*^+/−^),. **b-e,** Bar graphs showing the relative mRNA expression of *Parp6* mRNA in **b,** heart; **c,** lung; **d,** liver; **e,** Spleen in 6-7 months-old Flox and *Parp6*^+/−^ mice. **f-g,** Immunoblot for PARP6 in (**f**) lung and (**g**) liver lysates from Flox and *Parp6*^+/−^ mice. **h-k**, Echocardiographic assessment of cardiac performance in Flox and *Parp6*^+/−^ mice. Echocardiographic quantification of **(h)** Left ventricle posterior wall thickness at diastole (LVPWd); **i,** Left ventricle posterior wall thickness at systole (LVPWs); **j,** left ventricular internal diameter in diastole (LVIDd); and **k,** left ventricular internal diameter in systole (LVIDs); bars indicate mean with error as plotted. Statistical significance was assessed by unpaired two-tailed t-test or Man Whitney test for Flox and *Parp6*^+/−^ comparisons. Significance is annotated as ns (not significant), * = p < 0.05, ** = p < 0.01, *** = p < 0.001.

**Supplementary Fig. 2:**
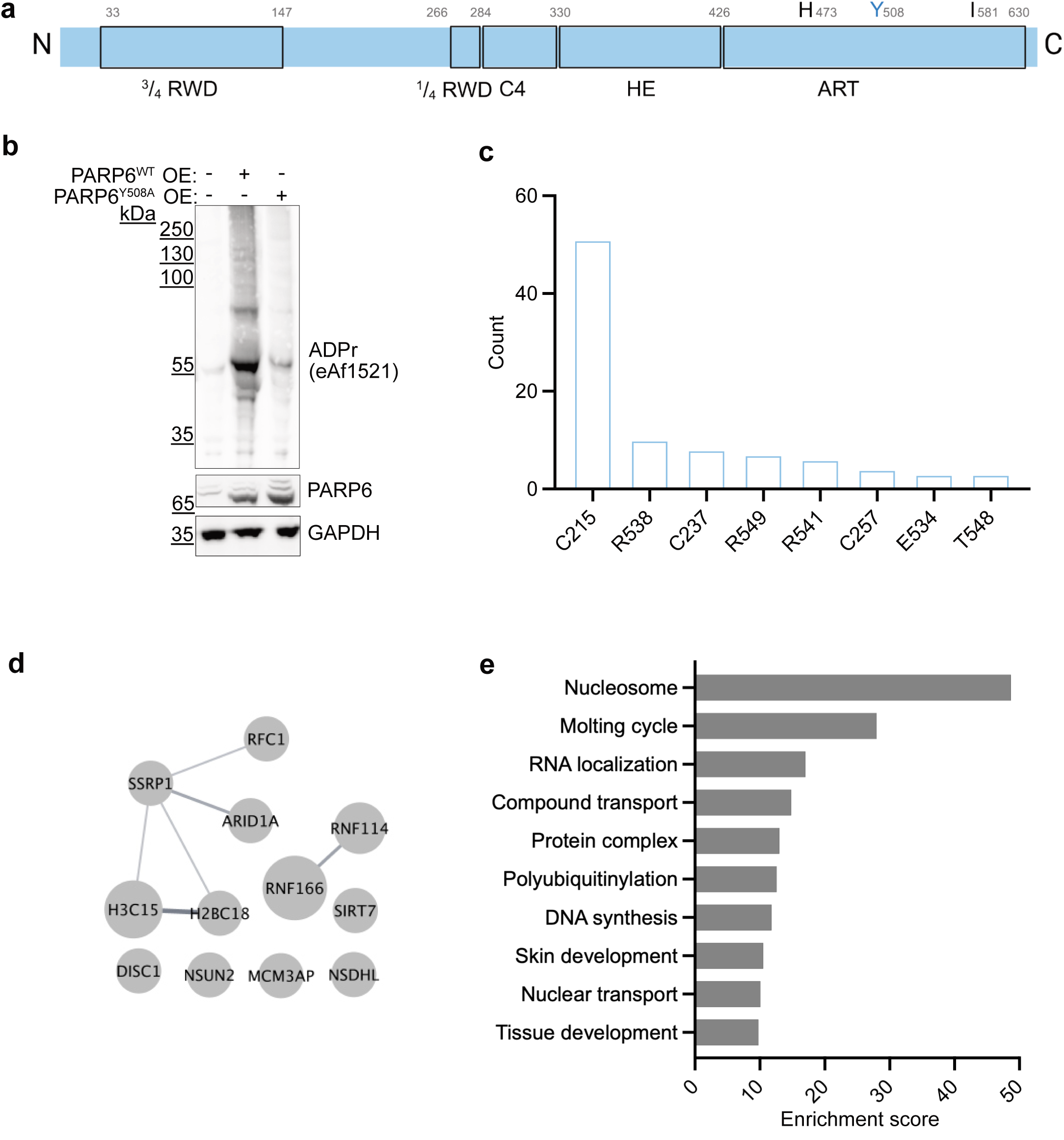
Characterization of PARP6^WT^ and PARP6^Y508A^. **a,** Schematic of PARP6 domains with domain structure (RWD: RWD domain, C4: central cysteine-rich C4 zinc-finger domain, HE: a helical subdomain, ART: catalytic domain) and catalytic triad residues indicated. **b,** Representative immunoblot depicting doxycycline-inducible (250 ng/ml) overexpression of PARP6^WT^ and PARP6^Y508A^ in AC16 cardiomyoblasts and corresponding ADPR levels. GAPDH is shown as a loading control (n=3). **c,** Bar graph depicting PARP6^WT^ automodification sites as PSM counts in AC16 cardiomyoblasts upon PARP6^WT^ overexpression conditions (induced by 250 ng/ml doxycycline). **d,** STRING interaction network of proteins with ≥1.5-fold higher ADPr spectral counts in PARP6^Y508A^ overexpression compared to PARP6^WT^ overexpression in AC16 cardiomyoblast cells; node size scaled with ADPr spectral counts. Unconnected nodes were omitted. **e,** Gene Ontology enrichment analysis of the proteins enriched in PARP6^Y508A^ overexpression (induced by 250 ng/ml doxycycline) **(d)** Gene Ontology-term enrichment for proteins with ≥1.5-fold higher ADPr spectral counts in PARP6^WT^ overexpression compared to PARP6^Y508A^ overexpression (induced by 250 ng/ml doxycycline). None of the terms were significant after multiple-comparison correction.

**Supplementary Fig. 3:**
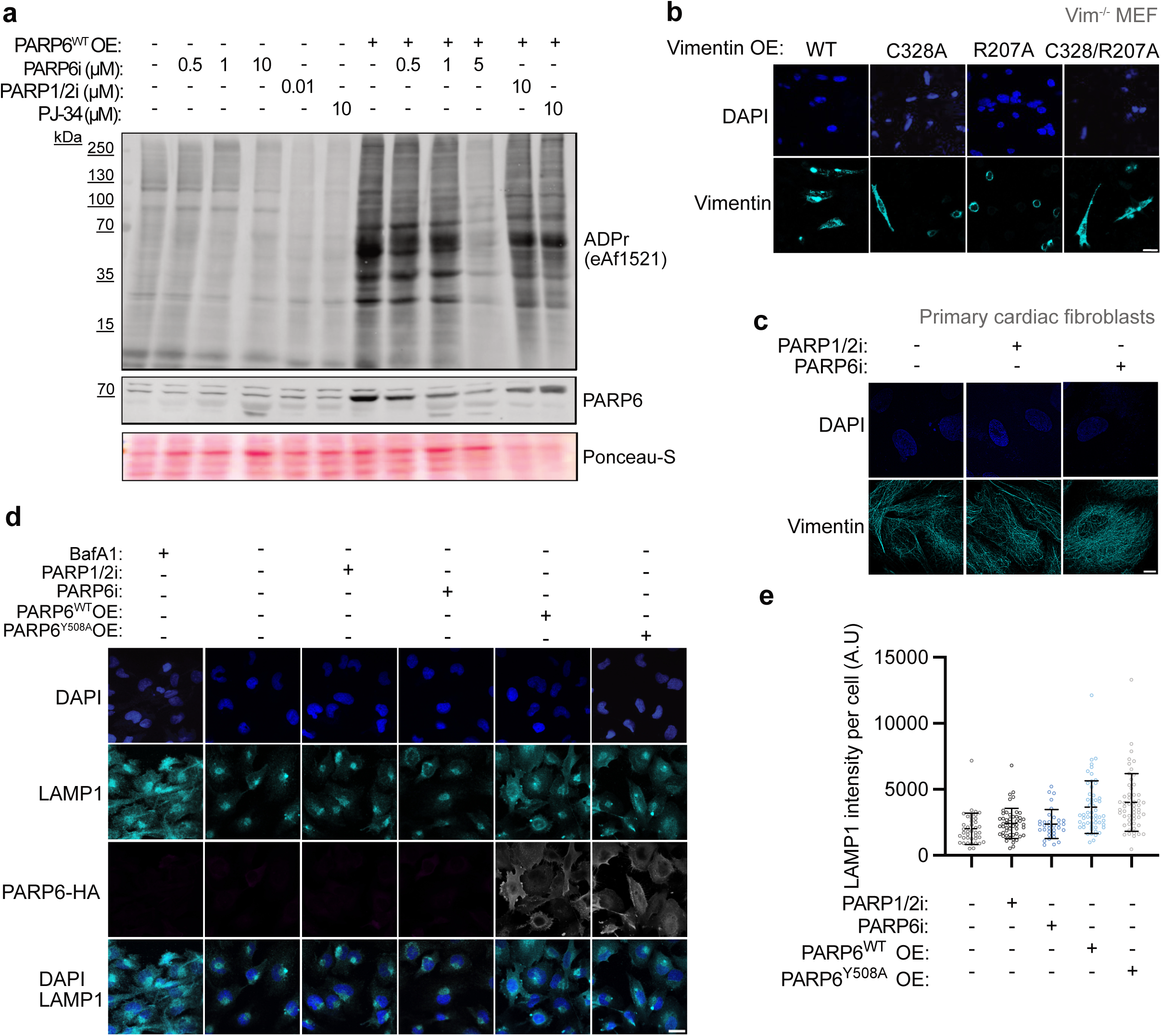
Vimentin filament formation and lysosomal positioning upon PARP6 inhibition. **a,** Representative immunoblot for ADP-ribosylation (ADPr) and PARP6 in PARP6^WT^ overexpressing AC16 cells treated with AZ0108 (PARP6i, indicated concentrations), PARP1/2i (indicated concentrations,) or PJ-34 (10 µM). Ponceau S staining is shown as the loading/transfer control, (n=2) **b,** Representative confocal microscopy images of Vimentin^−/−^ MEFs overexpressing Vimentin^WT^, Vimentin^C328A^, Vimentin^R207A^, or Vimentin^C328A/R207A^. Images show DAPI-stained nuclei (blue) and vimentin (cyan) (n=3). Scale bar represents 50 µm. **c,** Representative super-resolution microscopy images of primary cardiac fibroblasts treated with PARP1/2i (100 nM) or PARP6i (1 µM). Images show DAPI (blue) and vimentin (cyan) (n=2). Scale bar represents 20 µm. **d,** Representative confocal microscopy images of cells treated with PARP1/2i (100 nM) or PARP6i (1 µM), and/or PARP6 overexpression as indicated, and stained for LAMP1. The autophagy-inducer bafilomycin (BafA1, 100nM) was used as a positive control. Images show DAPI (blue), LAMP1 (cyan), and PARP6-HA (grey) (n=1). Scale bar represents 20 µm. **e,** Quantification of LAMP1 intensity from **(d)**. Violin plots show single-cell LAMP1 intensity per cell (A.U.) across the indicated treatments (n=1; 30 cells)

**Supplementary Fig. 4:**
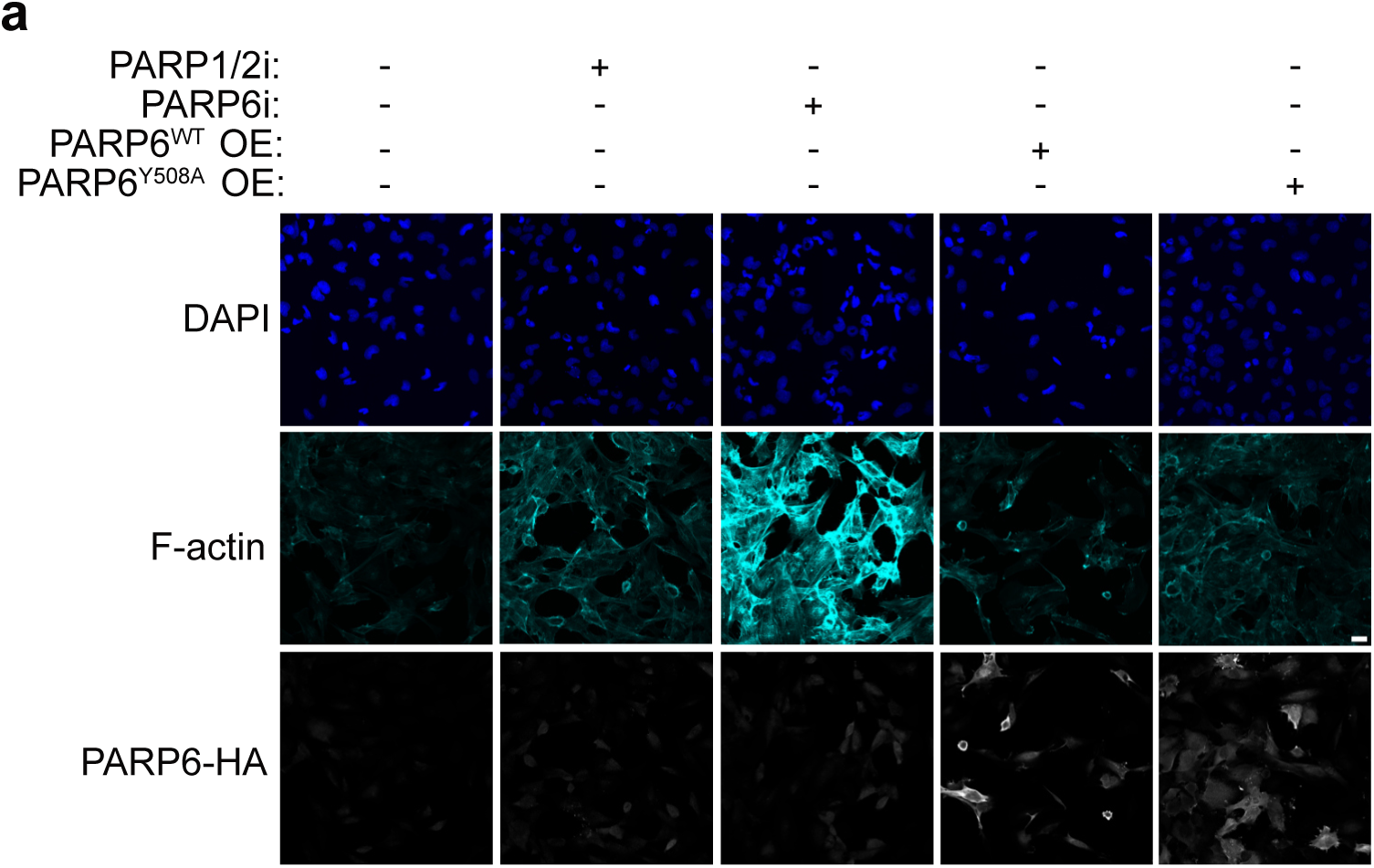
PARP6i treatment and actin stress fibers. **a,** Representative confocal microscopy images of AC16 cells treated with PARP1/2i (100 nM) or PARP6i (1 µM), overexpression of PARP6^WT,^ or PARP6^Y508A^ (induced by 250 ng/ml doxycycline). Images show DAPI (blue), F-actin (cyan), and PARP6 (anti-HA, yellow) (n=3). Scale bar represents 20 µm.

**Supplementary Fig. 5:**
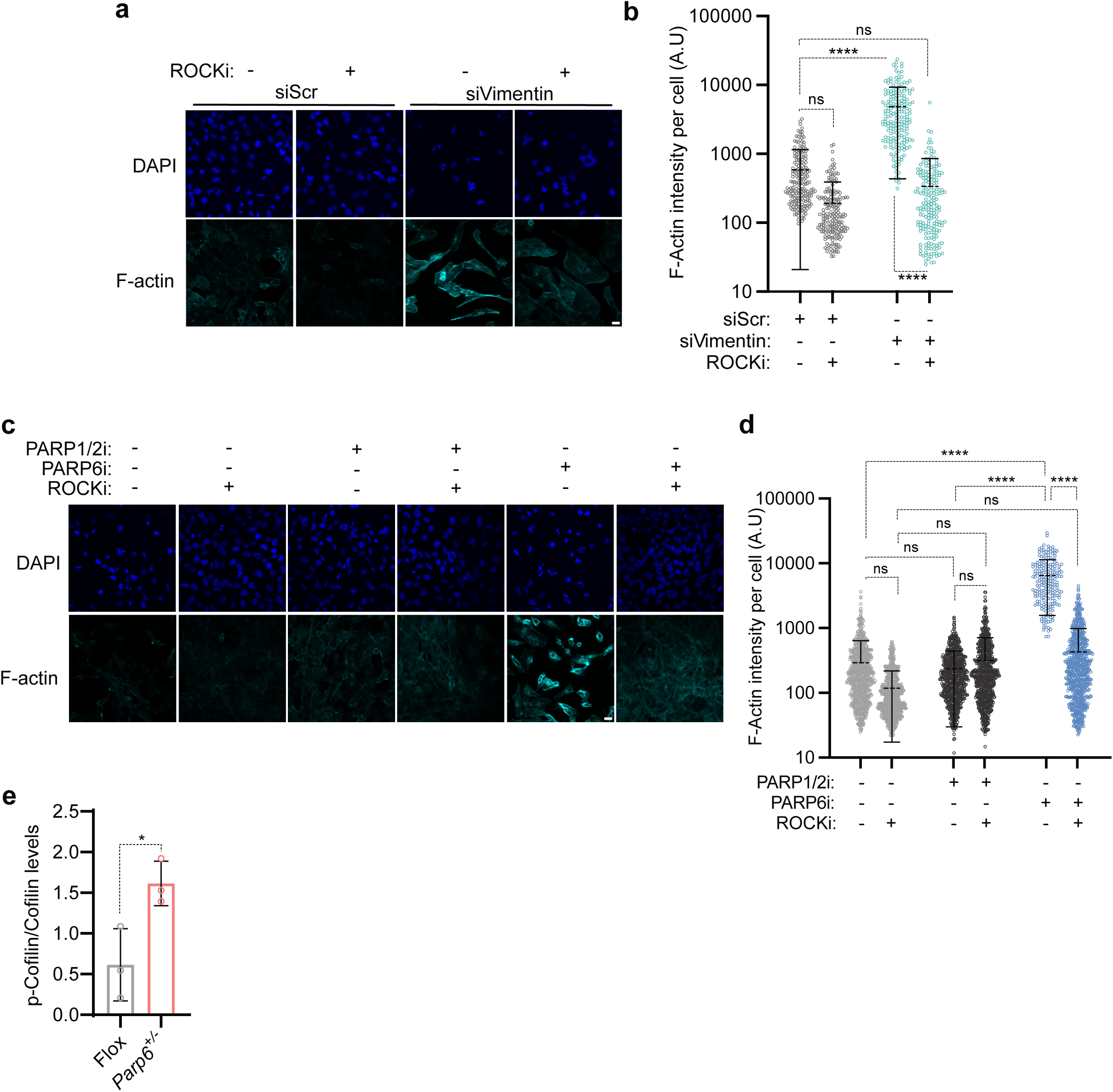
ROCK as a downstream effector of RhoA signaling. **a,** Representative confocal microscopy images of siScr and siVimentin-treated (10 nM) cells under ROCKi-treated (10 µM) conditions. Images show DAPI (blue) and F-actin (cyan) (n=3). Scale bar represents 20 µm. **b,** Scatter plot quantification of F-actin intensity per cell from **(a)** (n=3; at least 200 cells were quantified per condition). Statistical testing was performed on the means of 3 independent experiments using one-way ANOVA with multiple-comparison correction. **c,** Representative confocal microscopy images of AC16 cells treated with PARP1/2i (100 nM), PARP6i (1 µM), alone or in combination with ROCKi (10 µM). Images show DAPI (blue) and F-actin (cyan) (n=3). Scale bar represents 20 µm. **d,** Scatter plot quantification of F-actin intensity per cell from **(c)** (n=3; at least 200 cells were quantified per condition). Statistical testing was performed on the means of 3 independent experiments using one-way ANOVA with multiple-comparison correction. **e,** Bar plot representing the quantification of p-cofilin normalized to total cofilin levels in 6-7 months-old *Parp6^+/**−**^* hearts relative to age-matched Flox controls. ns = non-significant, * = p < 0.05, ** = p < 0.01, *** = p < 0.001, **** = p < 0.0001.

